# Passenger-surface microbiome interactions in the subway of Mexico City

**DOI:** 10.1101/2020.04.28.067041

**Authors:** Daniela Vargas-Robles, Carolina Gonzalez-Cedillo, Apolinar M. Hernandez, Luis D. Alcaraz, Mariana Peimbert

## Abstract

Interaction between hands and the environment permits the interchange of microorganisms. The Mexico City subway is used daily by millions of passengers that get in contact with its surfaces. In this study, we used 16S rRNA gene sequencing to characterize the microbiomes of frequently touched surfaces, also comparing regular and women-only wagons. We also explored the effect of surface cleaning on microbial resettling. Finally, we studied passenger behavior and characterized microbial changes after traveling.

Most passengers (99%), showed some type of surface interaction during a wagon trip, mostly with the hands (92%). We found microbiome differences associated with surfaces, probably reflecting diverse surface materials and usage frequency. The platform floor was the most bacterial diverse surface, while the stair handrail and pole were the least diverse ones. After pole cleaning, the resettling of microbial diversity was fast (5-30 minutes); however, it did not resemble the initial composition.

After traveling, passengers significantly increased their hand microbial diversity and converged to a similar microbial composition among passengers. Additionally, passenger hand microbiomes resembled subway surfaces in diversity and also in the frequency of potentially pathogenic taxa. However, microbial fingerprints were preserved within passengers after traveling.

## Introduction

Mexico City’s subway transports around 1,678 million passengers per year (4.2 million daily), making it the ninth largest transit subway in the world (UITP, 2018). This high number of visitors promotes multiple physical interactions, becoming an essential system for studying colonization and dissemination of microbes.

Hands are an important channel of the interactions of humans with their surroundings. The hand microbiome is mainly comprised of the phyla Actinobacteria, Proteobacteria, and Firmicutes (Gao et al., 2007). Hand microbiomes are highly diverse, surpassing oral and intestinal microbiomes (Costello et al., 2009). They highly vary among individuals, but also between the right and the left hand of the same person (Fierer et al., 2008). Constant exposure to diverse environmental sources and perturbations (e.g., handwashing) is a factor of the hand’s microbiome heterogeneity (Fierer et al., 2008).

Interaction between hands and the environment permits the interchange of microorganisms, explaining the high proportion of human-associated microbes in built environments (Lax et al., 2014). There is a human microbiome signal that is strong and traceable among people and buildings (Fierer et al., 2010). Cohabitation results in closer microbial composition than kinship, with more similar microbial profiles of people sharing the same house compared to others (Lax et al., 2014). Additionally, the surfaces in one house can be more similar in terms of microbiome composition when compared to different houses (Ruiz-Calderon *et al.*, 2016). Similarly, there are microbe differences in built areas used exclusively by women or men; for example, *Lactobacillus iners* was found as a female-associated bacterium, while *Dermabacter hominis, Facklamia,* and *Corynebacterium* were more abundant in rooms used by males (Luongo et al., 2017; Fierer et al., 2008; Ross, Doxey, & Neufeld, 2017; Takagi et al., 2019). There is evidence that indoor and human microbiomes are closely related and influence each other.

The Mexico City subway contains different microenvironments (Hernández-Castillo et al., 2019). Most train lines are devoid of sunlight, and only a few lines may run in the exterior a fraction of their route. Exterior air is ventilated into the subway from a ducted air stream, and indoor air is recirculated using exhaust fans. Particulate matter levels are higher inside stations than outdoors (Mugica-Álvarez et al., 2012). In the hot season, water is spread out into the air by fans. While the statin floors are cleaned daily, train wagons are deep-cleaned once a month. Other surfaces such as turnstiles, stairs, and escalator handrails are cleaned eventually with a not strict schedule.

The subway is the most used transportation system in Mexico City. Metro travelers in Mexico City face daily peculiarities such as the sale and consumption of food inside the premises, street vendors, beggars, and the absence of seats at the stations. The first two train cars are exclusively for women, disabled, and the elderly (Dunckel-Graglia 2013). Some cities have applied exclusive cars for women as a measure to decrease sexual harassment (Junior et al., 2017; Horii et al., 2012).

There are culture-independent studies of subway surfaces of New York City (Afshinnekoo et al., 2015), Boston (Hsu et al., 2016), Oslo (Gohli et al. 2019), Mexico City (Hernández et al. 2019), and by MetaSUB (MetaSUB International Consortium, 2016), an international initiative. Such studies have shown that the subway microbiome is structured mostly by commensal bacteria from the skin and that microbial composition and diversity vary according to the material and type of usage. In the Hong Kong subway, there are differences between morning and afternoon microbial composition, with more antibiotic resistance genes in the afternoon (Kang et al., 2018). In the same study, the commuters’ hand microbiomes were explored measuring bacteria acquisition in a 30-minute trip (Kang et al., 2018).

The present study describes the interaction between the Mexico City subway microbiota and those of its passengers. We compared microbiomes from different subway surfaces comprising stations and regular and women-only wagons. We also evaluated the velocity of the bacterial succession after an event of surface cleaning. Additionally, we characterized the passengers’ microbiomes before and after traveling (Fig. 1, Table S1).

**Figure 1.**
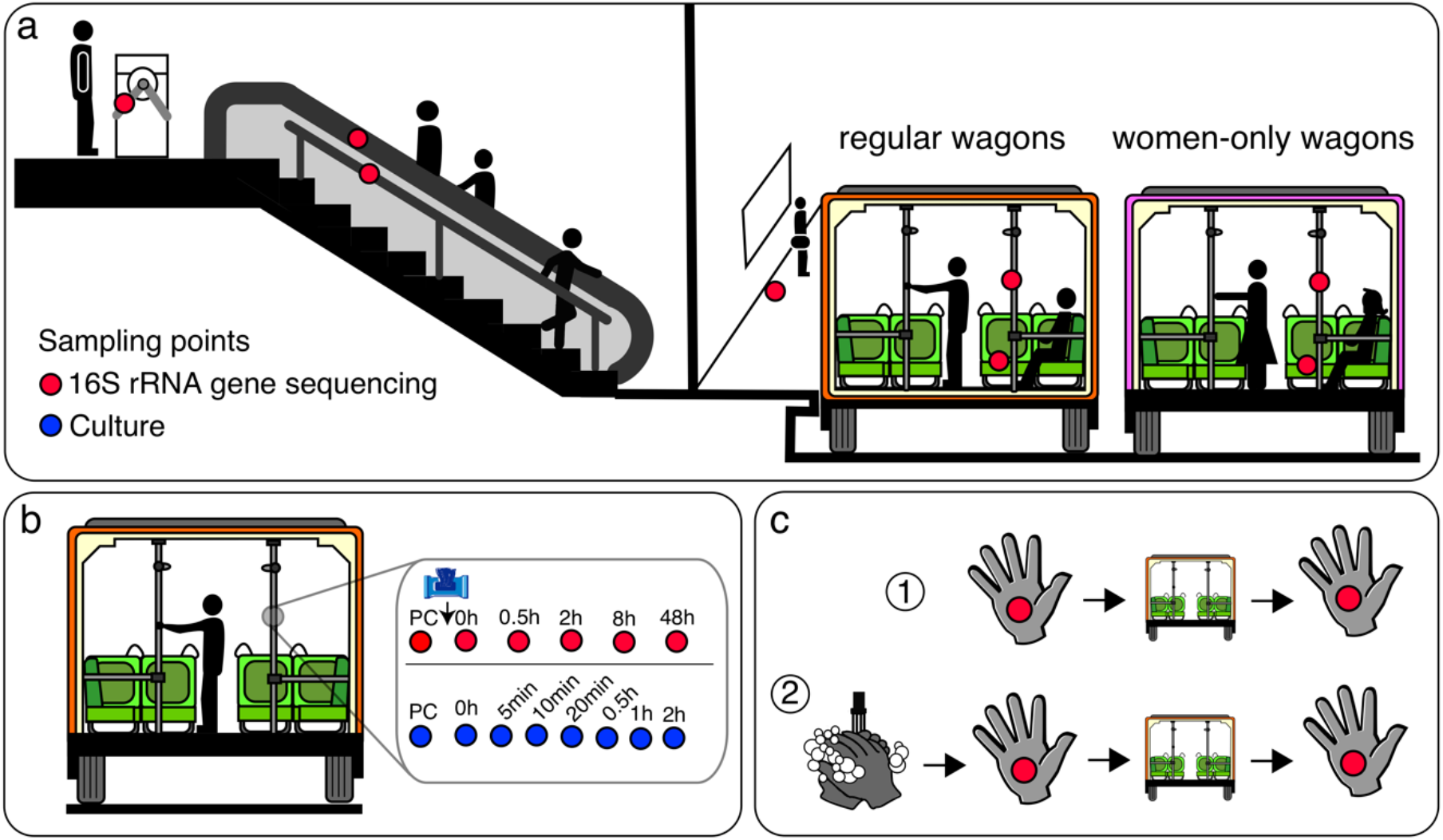
Study design. We swabbed surfaces and used the 16S rRNA gene (red dots) and microbial cultures (blue dots) to describe bacterial diversity. (a) Microbiome comparison of subway surfaces: turnstiles, escalator handrails, stair handrails, platform floors, poles, and train seats. Poles and train seats were sampled in regular and women-only wagons (N = 5 per site). (b) Microbiome succession study after a cleaning event in poles. Microbiome changes were evaluated at pre-cleaning (PC) and at 0, 0.5, 2, 8, and 48 h after cleaning. (c) Hand microbial diversity before and after traveling; we evaluated the effect of traveling with and without previous handwashing.

**Table S1.**
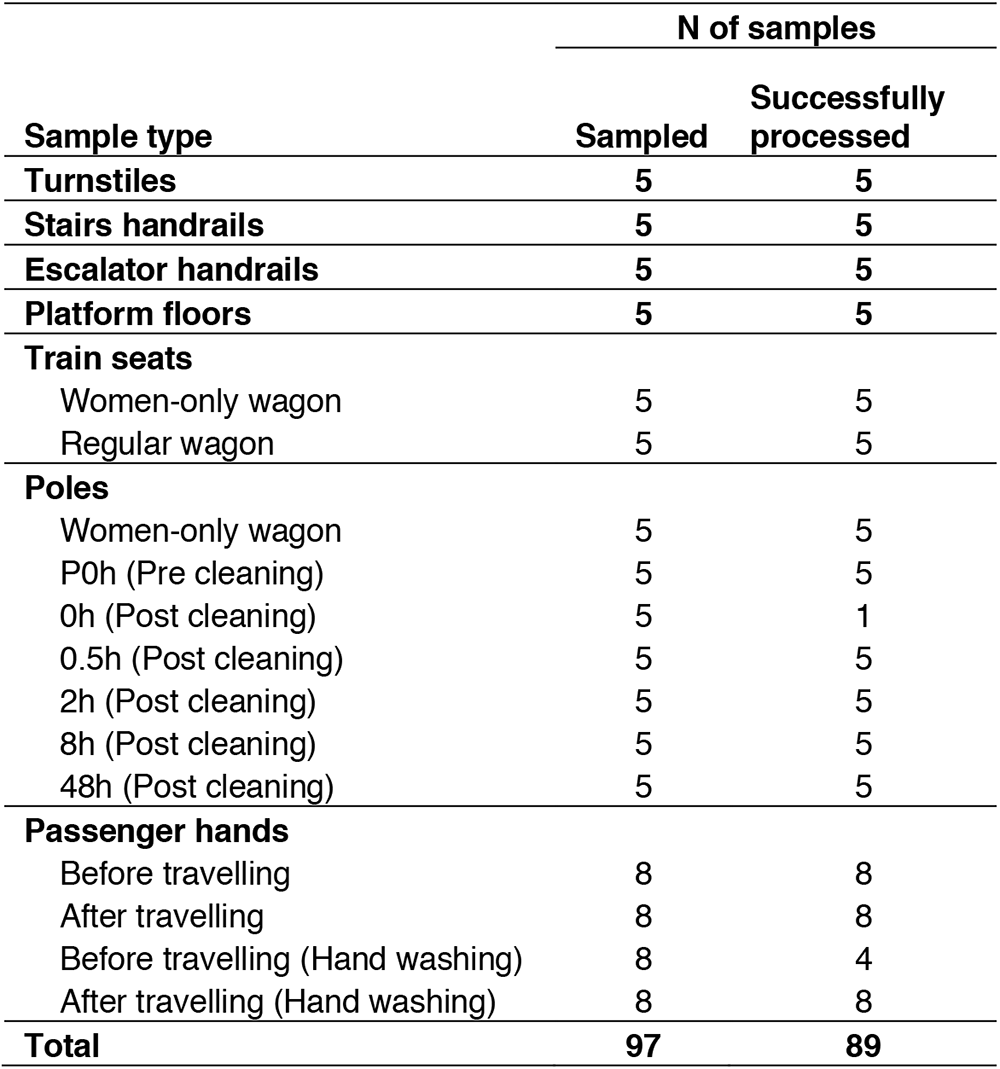
Number of samples collected and successfully processed.

## Results

### Passenger behavior and subway interaction

To understand the type and level of interaction that a passenger has with the subway environment, we traveled with passengers and registered their behavior during eight different weekdays. A total of 120 passengers were randomly picked and observed during a train trip (67 adults and 53 elders), from the entrance to the end of their trips (Table S2). Prevalence (%) and median frequency (times/10 min) of contact with particular surfaces or objects were registered. Most of the passengers (99%) had contact with any wagon surface; 92% of the contacts were with the hands and the rest with the body or cloth. As expected, the hands were the most common means of interaction with train surfaces (4.1 times/10 min), with higher frequency in the older people than in adults (5.0 vs. 3.6 times/10 min; *p* = 0.030, Wilcoxon test). Most users touched train poles (89%), being the most recurrent touched surface (marginally higher in the elderly than in adults, 4.9 vs. 4.1 times/10 min; *p* = 0.051, Wilcoxon test). Self-contact of the passengers was measured, and the face/head was the most commonly touched body part (73%, 2.2 times/10 min), similar in frequency between age groups. In face/head area touching, the skin predominated (68%, 1.3 times/10 min), followed by the hair/scalp (27%, median of 0, mean of 0.7 times/10 min), any mucosa (24%, median of 0, mean of 0.2 times/10 min), and the ear canal (6%, median of 0, mean of 0.04 times/10 min), similar between age groups. Passenger hands were also in frequent contact with personal articles (73%, 1.7 times/10 min), with cell phones being the most commonly and frequently used items in adults compared to the elderly (49 vs. 5.7%, median of 0 for both groups and mean of 0.9 vs. 0.05 times/10 min respectively; *p* = 3 x 10^-7^, Wilcoxon test). Other activities not directly related to hands, such as sitting, were also higher in the elderly (*p* = 0.001, Wilcoxon test), as well as laying the body on any other surface than seats (*p* = 0.049, Wilcoxon test). Further activities with microbiological relevance were also registered but observed to a lesser extent, such as touching other people, money interchange (buying or charity), coughing, drinking, eating, and eating with bare hands. Putting on makeup, book reading, and sleeping were also eventually observed, although not included in the table.

**Table S2:**
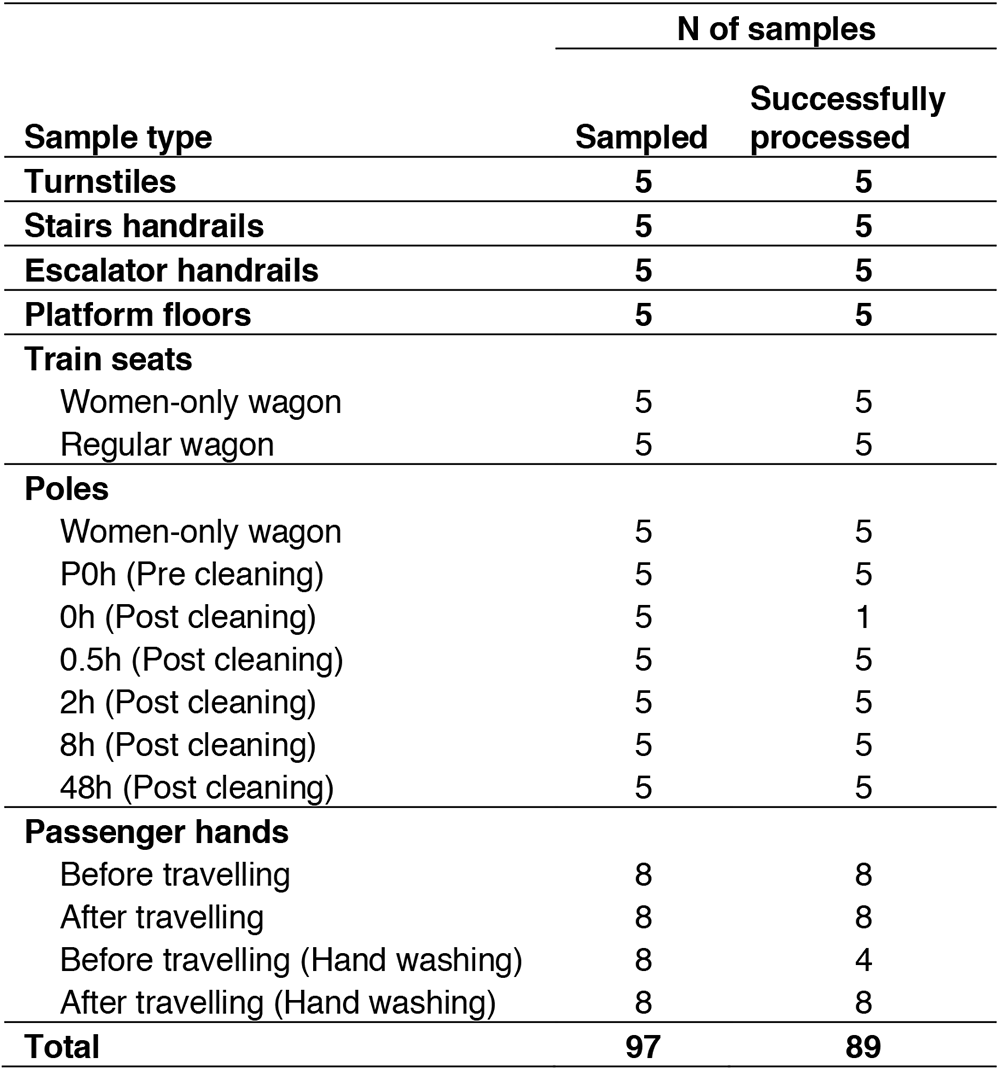
Frequency and prevalence of activities observed in passengers during a train trip.

**Table S3:**
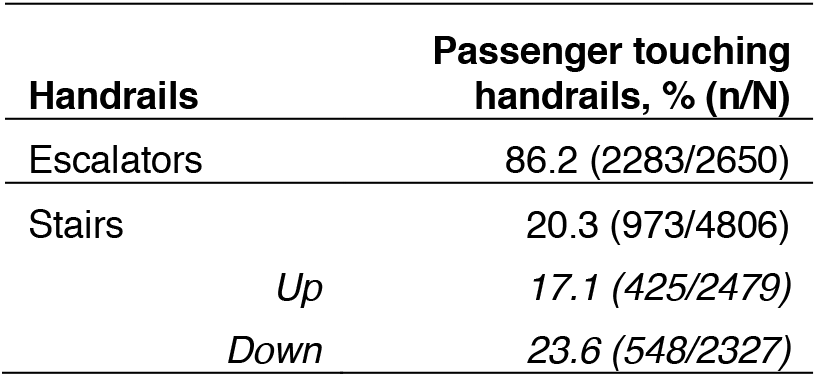
Percentage of passengers touching the handrails of escalators and stairs.

Additionally, we observed passengers using the escalator and stairs in the stations (N = 7,456 passengers, Table S3). Escalator handrails were touched continuously with the hands (86.2%, N = 2,650), while stair handrails were less frequently used (20.3%, N = 4,806), being higher for stairs going down than going up (23.6% vs. 17.1% respectively, *p* < 1 x 10^-7^, Chi-squared test).

### Massive sequencing of 16S rRNA gene

We sequenced a total of 89 samples, with 10,538,220 reads being obtained, which resulted in 5,238,317 paired sequences of an average length of 450 bp (Tables S4 and S5). Sequences were filtered to discard singleton, chloroplast, or mitochondrial sequences. With an average of 34,125 sequences per sample, we identified a total of 75,914 97% OTUs (1,121 genera). All samples were rarefied at the minimum sequence number per sample (6,242 sequences). Subsampling generated 29,811 OTUs (939 genera; Table S5). A rarefied data set was used to present the results of this study. We did not identify archaeal OTUs, and only 28 OTUs (0.004% of the sequences) did not match with any known organism.

**Table S5.**
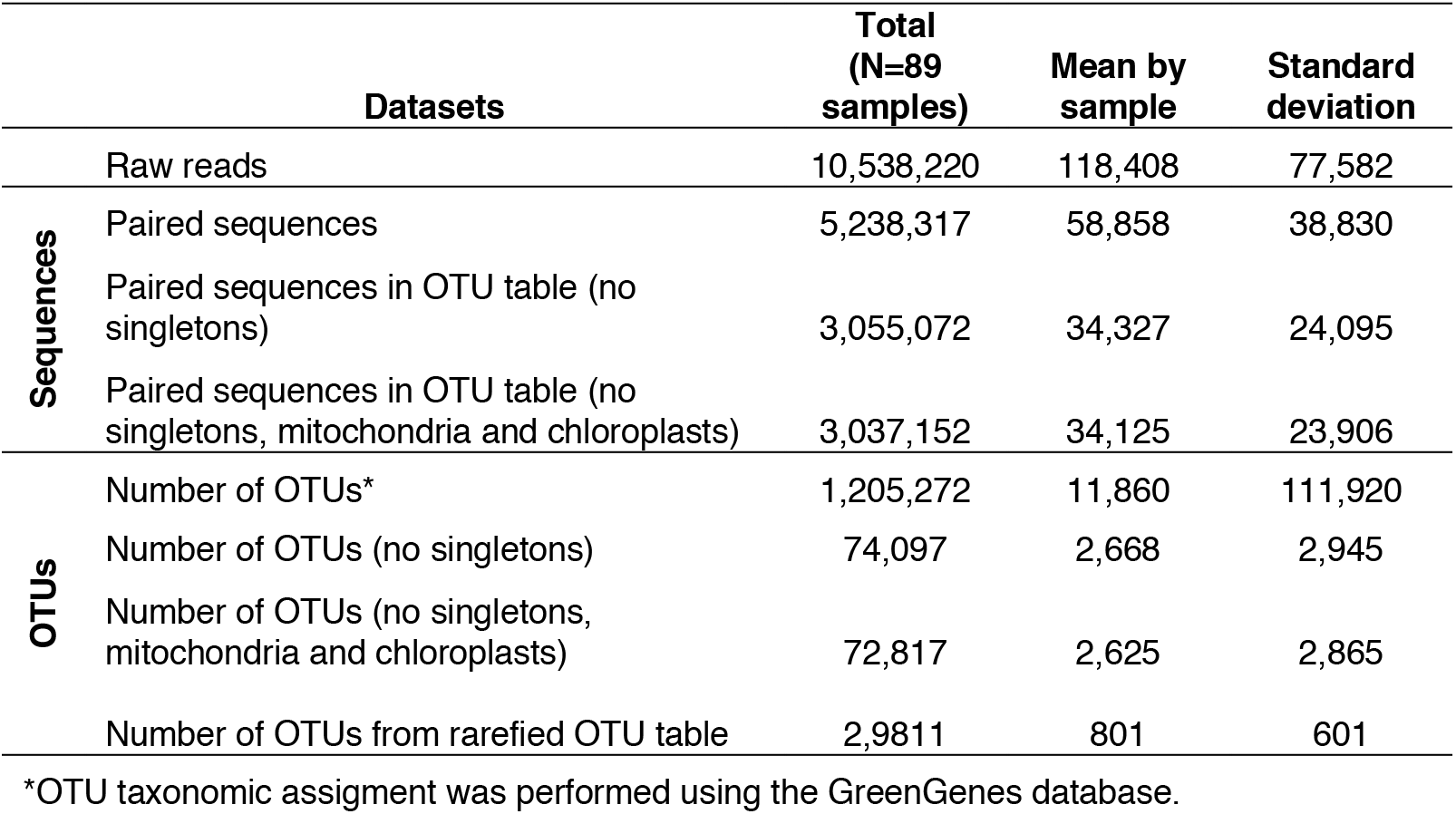
Numbers of raw reads, paired-end reads, and OTUs.

### Subway surface microbiome

We sampled surfaces from turnstiles, escalator handrails, stair handrails, platform floors, poles, and train seats (Fig 1a). Relative abundances of surfaces microbiomes were higher for the phyla Proteobacteria (31%), Actinobacteria (30%), Firmicutes (24%), Bacteroidetes (9%), and Fusobacteria (1.5%). At the genus level, the five most abundant taxa comprised 36% of the total abundance: *Acinetobacter* (10.5%), *Corynebacterium* (8.2%), *Streptococcus* (7.3%), *Staphylococcus* (6.8%), and *Propionibacterium* (3.4%). Only nine genera were ubiquitous across all 40 samples (Table S6), comprising 44% of the overall abundance. Figure 2a shows a summary of the genera composition. The five most abundant OTUs were *Acinetobacter* sp. (5.6%), *Staphylococcus* sp. (3.8%), *Propionibacterium acnes* (3.6%), *Streptococcus* sp. (2.7%), and *Staphylococcus epidermidis* (2.0%).

**Figure 2.**
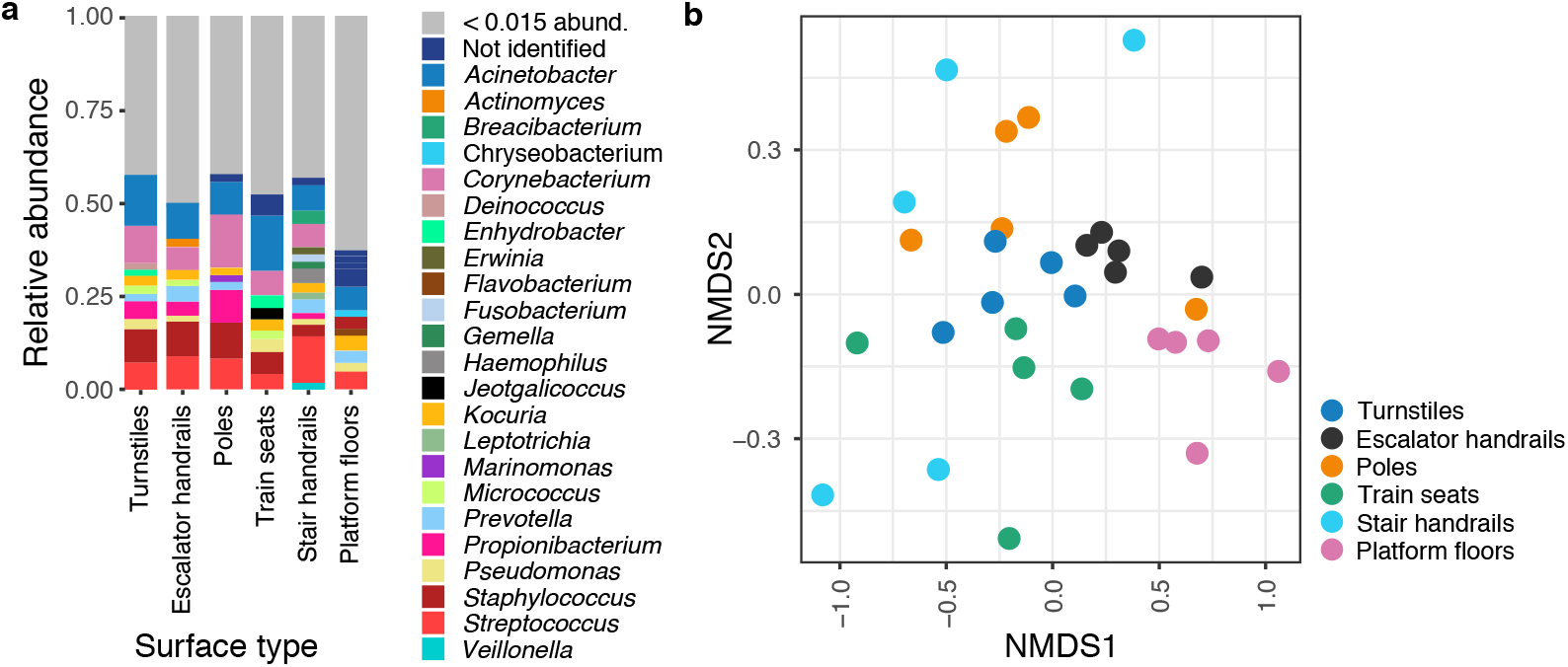
Subway surface microbiome diversity. Microbial composition differed among subway surface types. (a) Taxa summary showing the most abundant genera. (b) Beta diversity at the genus level, Bray-Curtis based non-metric multidimensional scaling (NMDS) plot of surface types samples.

**Table S6.** Shared genera among samples within the same surface type and among all samples.

| Surface types | N | Shared genera |
| --- | --- | --- |
| Platform floors | 5 | 148 |
| Escalator handrails | 5 | 138 |
| Stair handrails | 5 | 26 |
| Turnstiles | 5 | 66 |
| Poles (regular wagons) | 5 | 49 |
| Poles (women-only wagons) | 5 | 17 |
| Train seats (mix wagons) | 5 | 39 |
| Train seats (women-only wagons) | 5 | 28 |
| Total shared taxa among all samples | 40 | 9* |

\**Acinetobacter*, *Corynebacterium*, *Streptococcus*, *Staphylococcus*, *Propionibacterium*, *Kocuria*, *Pseudomonas*, *Micrococcaceae*, *Micrococcus*

Alpha diversity varied among different surface types (for Shannon, Observed OTUs, Chao1, and Simpson metrics, *p* < 0.02, Kruskal-Wallis; Fig. S1 a). Platform floor was the most diverse surface, while the stair handrail and pole were the least diverse ones (*p* < 0.010, Dunn test; Fig. S1a). Microbial composition differed among surface types (*p* = 0.001, F = 1.99, PERMANOVA), with different variance dispersion (*p* = 0.018, F = 3.35, PERMADISP2; Fig. S1b).

Based on hierarchical clustering, the platform floor showed the most distinctive bacterial composition. Turnstiles, escalator handrails, and poles showed greater similarities (Fig. S1b and S1c). Interestingly, stair handrail samples did not cluster with other hand-contact surfaces, although they displayed the highest variance dispersion, reflecting high heterogeneity among samples. In contrast, escalator handrails and turnstiles showed the lowest dispersion, reflecting higher homogeneity among samples (Tukey’s HSD, *p.adjust* < 0.026, Fig. S1d).

We also explored microbiome differences between regular and women-only wagons for poles and seats and found no differences for any surface in terms of alpha or beta diversities (Fig S2). Further analysis was performed using amplicon sequence variants (ASVs), also without differences (see below).

**Figure S1.**
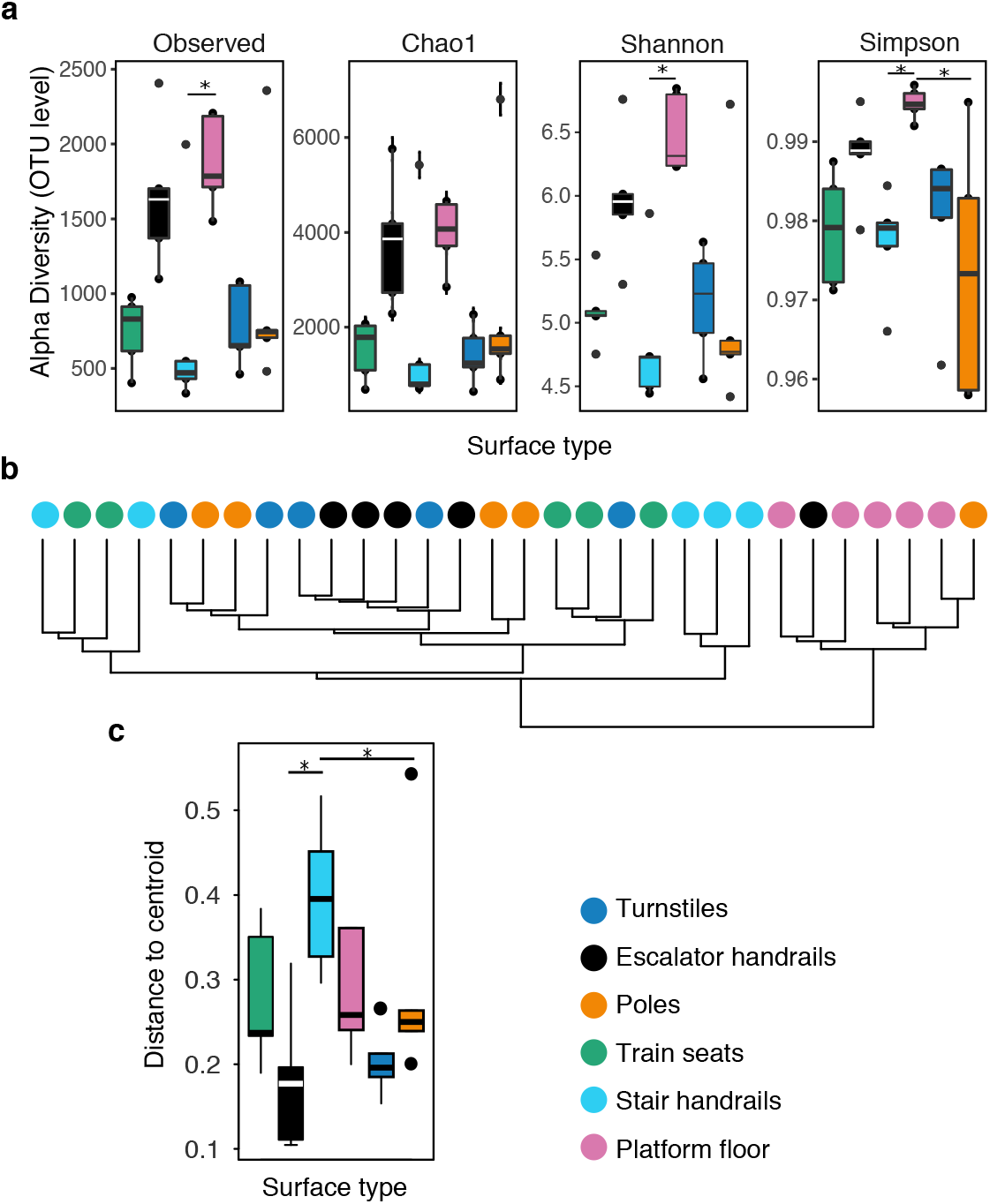
Subway surface microbiome diversity in stations and regular train wagons (N = 5 samples per surface type). The platform floor was most diverse, with the most distinctive composition. (a) Alpha diversity measures for surface type at the OTU level. Pairwise comparison showed that platform floor diversity was higher than that of stair handrails and poles (* *p* < 0.01, Nemenyi-tests). (b) Hierarchical clustering of individual surface samples, colored by surface type. Hierarchical clustering analyses were performed with the ward.2 method and Bray Curtis dissimilarity. (c) Variance dispersion among surface types (* *p* < 0.02, PERMADISP2; *p.adjust* < 0.026, Tukey’s HSD). Distances to centroid groups were calculated by reducing the original Bray Curtis dissimilarity to principal coordinates.

**Figure S2:**
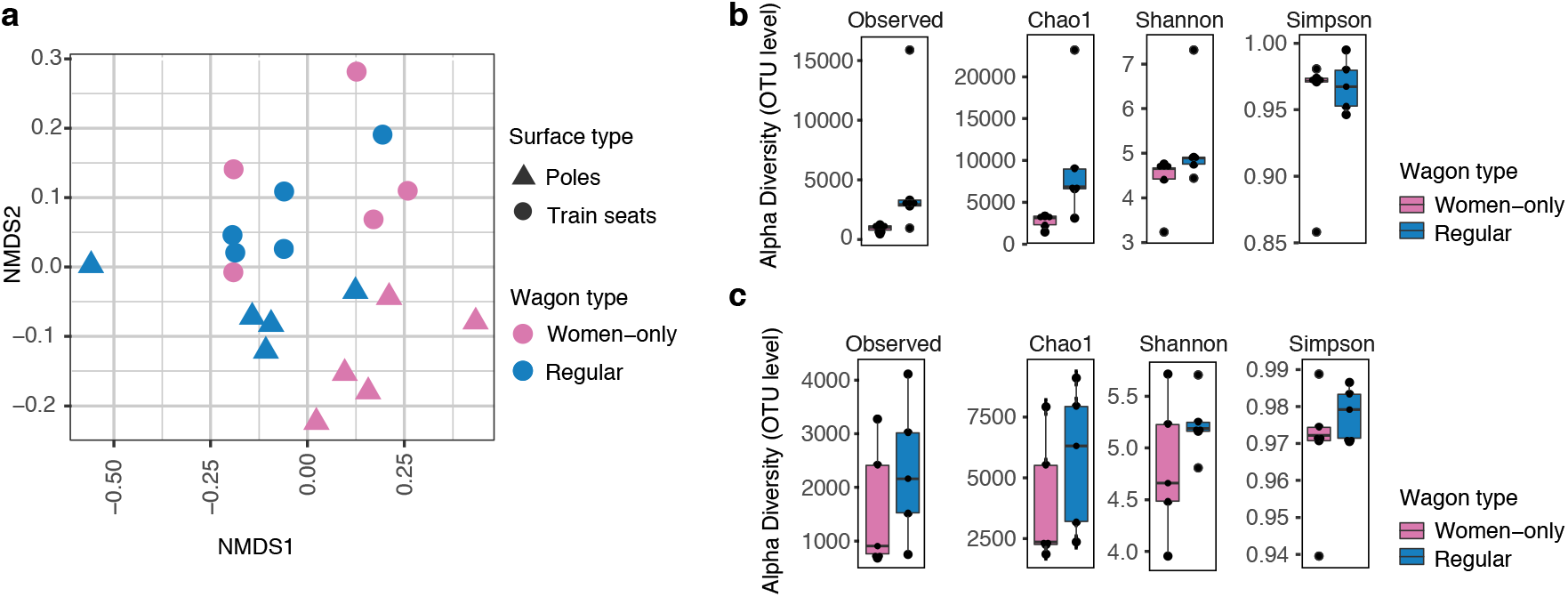
There are no microbiome differences between regular and women-only train wagons at the OTU level (N = 5 samples per category). (a) Non-metric multidimensional scaling (NMDS) ordination with Bray dissimilarity showing spatial distribution of sample groups (poles, *p* > 0.18, F = 1.13; train seats, *p* > 0.59, F = 0.95, PERMANOVA; NMDS stress = 0.20). (b) Alpha diversity measures at the OTU level. No significance was found between groups for any measure (*p* > 0.5, Kruskal-Wallis).

### Microbiome ecological succession after surface cleaning

We explored the changes in surface microbiomes after a cleaning event. We cleaned five poles with disinfectant towels and distilled water, and the poles were sampled pre-cleaning (PC) and along five time-points post-cleaning (0, 0.5, 2, 8, and 48 h) (Fig. 1b).

Cleaning of poles significantly reduced sample biomass, hindering 16S gene amplification. We only obtained 1/5 of PCR amplicons for the first time-point. Richness comparison among time groups (not including 0 h, N = 1) yielded significant results (Chao1, *p* = 0.038, Kruskal-Wallis). However, pairwise comparison removed this significance (*p* > 0.05, Dunn’s test; Fig. 3a). Beta diversity showed significant differences among group compositions (*p* = 0.001, F = 1.46, PERMANOVA; Fig. 3b), while similar dispersion was observed (*p* = 0.867, F = 0.33, PERMADISP2). Set analysis suggests that after cleaning, pole microbiomes acquire a composition different to that of the pre-cleaning samples.

**Figure 3.**
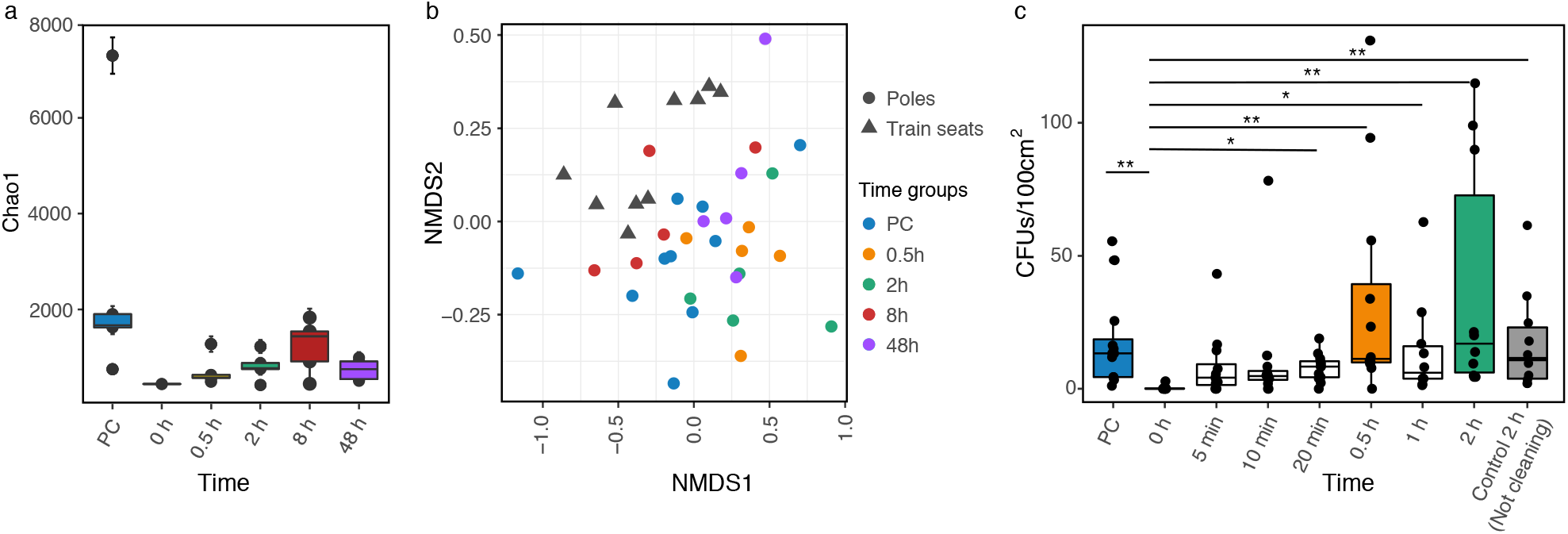
Pole microbiome diversity succession measured by 16S amplicons and colony-forming units (CFUs) after a cleaning event. (a) Alpha diversity and (b) beta diversity did not resettle within 48 h after cleaning. Nevertheless, the CFU count was regained within minutes (c). (a) Alpha diversity boxplots show the Chao1 richness estimator. (b) Non-metric multidimensional scaling (NMDS) ordination with Bray-Curtis dissimilarity at the genus level. (c) CFUs in seven time-points plus two controls: pre-cleaning (PC) and time control at 2 h (not cleaning). The last control pretends to evaluate natural changes of the microbiome in real time (\**p* < 0.05 and \*\**p* < 0.005, Dunn’s test).

The cleaning procedure removed 96.38% (3,867 OTUs) of the initial OTUs. A total of 67 OTUs (31 genera) were shared among all time groups, and they were not removed; these taxa included the most abundant genera. A total of 987 OTUs (234 genera) resettled the surfaces in at least one sample (Figs. 4 and S3). Many of them were intermittently identified, and only 422 initial OTUs (148 genera) were detected in the 48-h samples. Within 30 min, 369 removed OTUs (100 genera) resettled on the pole surfaces. The pre-cleaning group showed the highest count of unique taxa (Figs. 4 and S3), suggesting that rare taxa have not been completely established within 48 h. However, the high count of unique taxa in each time group, together with the intermittent identification of taxa, suggests that rare taxa are not persistent.

**Figure 4:**
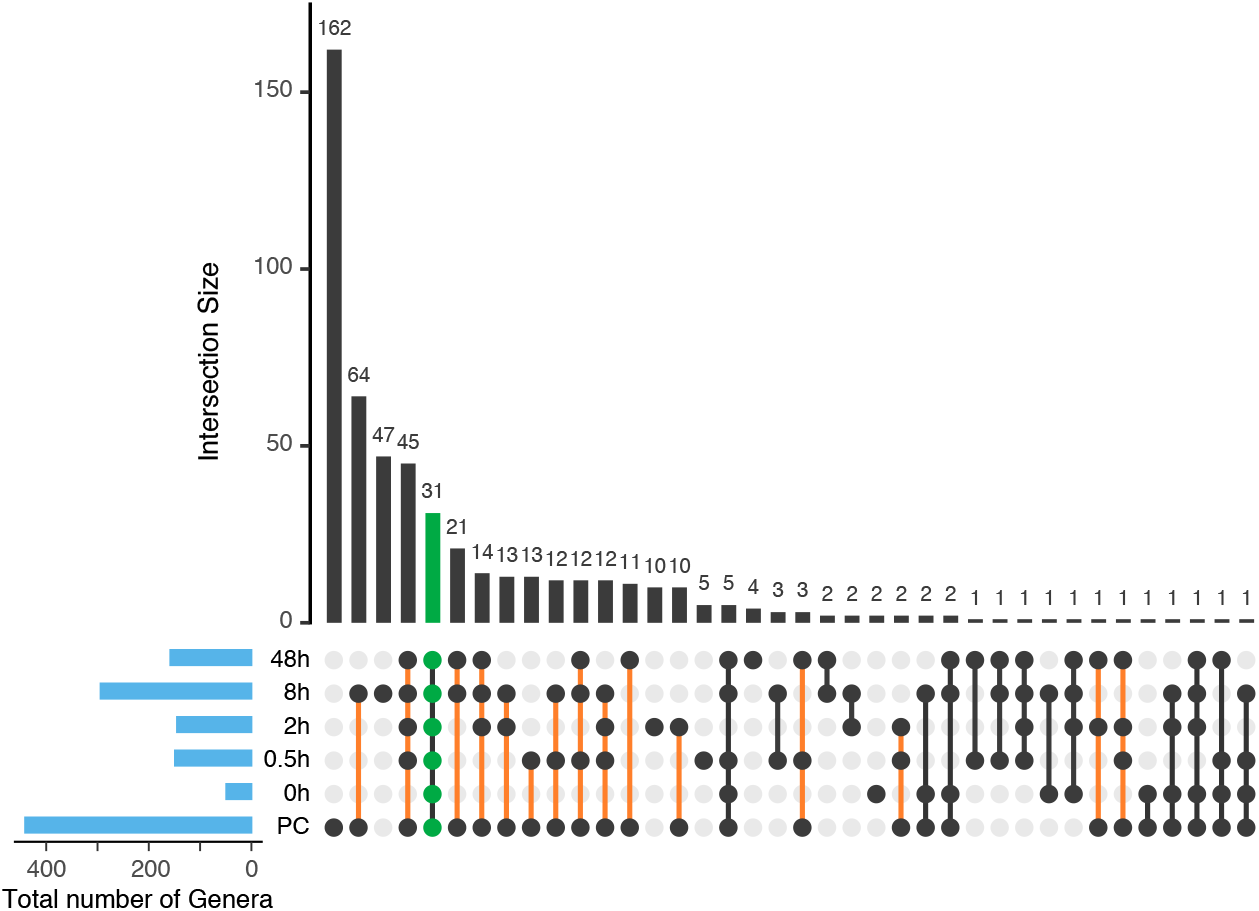
Unique and shared genera among time groups. Many genera are unique to each time group, and many resettled genera are not persistent. Upset plot of intersected genera among time points after cleaning; empty intersections are not shown. Taxa shared among all times are shown in green. Resettled taxa are shown with an orange line.

We also explored the dynamics of bacterial colonization for shorter periods (< 0.5 h) with a cultivation based-method. After the same cleaning procedure, 10-12 poles were sampled in seven time points: 0 h, 5 min, 10 min, 20 min, 0.5 h, 1 h, and 2 h. Results showed that 5 minutes were sufficient to reach a similar number of colony-forming units (CFU) when compared to the pre-cleaning group (PC vs. 5 min, *p* > 0.050, Dunn’s test; Fig. 3c).

**Figure S3:**
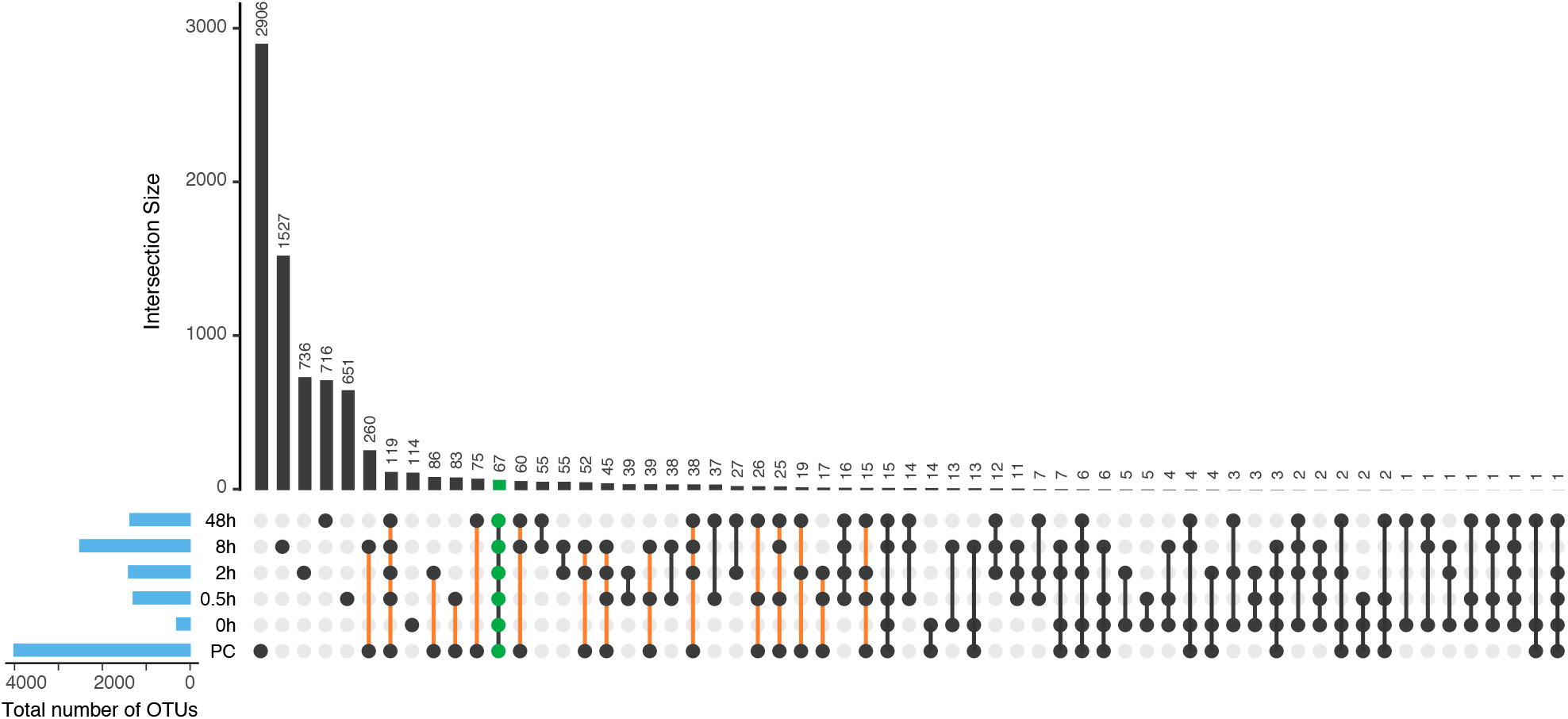
Unique and shared OTUs among time groups. Many OTUs are unique to each time group, and many resettled taxa are not persistent (a) Upset plot of intersected OTUs among time points after cleaning; empty intersections are not shown. Taxa shared among all times are shown in green. Resettled taxa are shown with an orange line.

### Passenger hand microbiome after traveling

Microbiome changes during a subway trip were measured in eight volunteers after an 11-station ride. Two procedures were evaluated: 1) traveling without previous handwashing and 2) traveling with previous handwashing (Fig 1c). Here, previous handwashing aimed to remove each passenger’s microbial signature and therefore to observe the newly acquired microbiota.

Subway traveling increased bacterial alpha diversity (OTU level: *p* < 0.020, for Observed, Chao1, and Shannon; Figs. 5a, S4a). The increased bacterial diversity may not be related with the touched surface type or the number of touched surfaces (*p* = 0.317, R^2^: 0.165; linear regression from net Shannon diversities and number of touched surfaces). Further exploration about the frequency and nature of the passenger-surface contact (intermittent or dragging-like) might help to elucidate this relationship.

**Figure 5.**
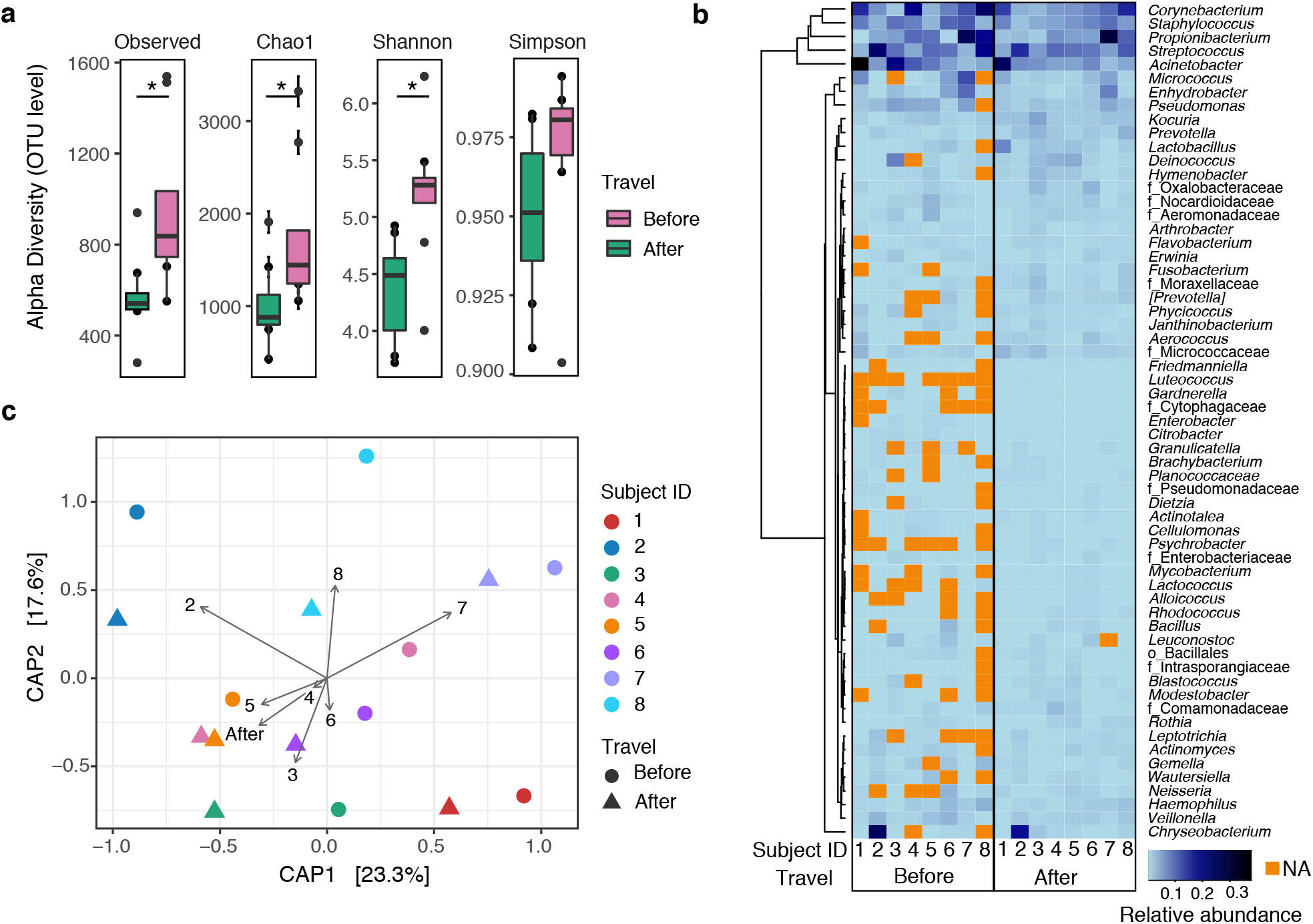
Changes in the passenger hand microbiome before and after traveling without handwashing. (a) Alpha diversity is increased after traveling (* *p* < 0.020). (b) Heatmap, using Manhattan distances, showing the relative abundance of all genera shared among subjects (columns, denoted by a number) before or after traveling. The subjects showed a closer microbiome profile after traveling. Letters before taxa indicate the best possible phylogenetic assignment (o: order and f: family). (c) Constrained analysis of principal coordinates (CAP) at the genus level, showing significant segregation for SubjectID and Travel (before and after) variables *(p* = 0.001, F = 2.4 and *p* = 0.006, F = 2.0, ANOVA-like permutation test for CAP).

After traveling, the observed OTUs increased by a mean count of 167% without (Table S7) and 408% with the handwashing procedure (Table S8). Unwashed hands lost 68% and washed hands 65% of their OTUs and acquired 135 and 254% of new OTUs, respectively. Unwashed hands conserved only 19% of the OTUs, while washed hands retained 35% of OTUs, which includes the most abundant ones. Additionally, small decreases of the most abundant taxa were observed after traveling with unwashed hands: *Acinetobacter* (11.7-7.7%), *Corynebacterium* (11.1-8.0%)*, Streptococcus* (10.2-8.7%), *Propionibacterium* (9.4-7.8%), and *Staphylococcus* (6.9-5.9%) (Fig. S5a). Similar changes were observed for most taxa for washed hands.

**Table S7:** Number of OTUs kept, lost, and newly acquired after one subway travel per passenger without handwashing.

|  | Poles | Escalator handrails | Stair handrails | Turnstiles | Train seats | Platform floors | Subject ID | N of OTUs |  |  |  |  | Shannon |  |
| --- | --- | --- | --- | --- | --- | --- | --- | --- | --- | --- | --- | --- | --- | --- |
|  |  |  |  |  |  |  |  | Before | After | Kept | Lost | Newly acquired | Before | After |
|  |  |  |  |  |  |  | 1 | 675 | 759 | 190 | 485 | 569 | 4.3 | 4.9 |
|  |  |  |  |  |  |  | 2 | 940 | 1539 | 326 | 614 | 1213 | 4.6 | 5.8 |
|  |  |  |  |  |  |  | 3 | 516 | 1512 | 197 | 319 | 1315 | 4.6 | 6.5 |
|  |  |  |  |  |  |  | 4 | 508 | 864 | 159 | 349 | 705 | 4.7 | 5.4 |
|  |  |  |  |  |  |  | 5 | 535 | 809 | 162 | 373 | 647 | 5.0 | 5.4 |
|  |  |  |  |  |  |  | 6 | 547 | 704 | 162 | 385 | 542 | 4.9 | 5.4 |
|  |  |  |  |  |  |  | 7 | 556 | 551 | 148 | 408 | 403 | 3.8 | 4.1 |
|  |  |  |  |  |  |  | 8 | 281 | 875 | 110 | 171 | 765 | 3.8 | 5.4 |
| Touched surface |  |  |  |  |  |  | mean | 569.8 | 951.6 | 181.8 | 388.0 | 769.9 | 4.5 | 5.4 |
|  |  |  |  |  |  |  | sd | 185.2 | 368.8 | 64.0 | 127.9 | 325.1 | 0.5 | 0.7 |

**Table S8:** Number of OTUs kept, lost, and newly acquired after one subway travel per passenger with handwashing.

| Poles | Escalator handrails | Stair handrails | Turnstiles | Train seats | Platform floors | Subject ID | N of OTUs |  |  |  |  | Shannon |  |
| --- | --- | --- | --- | --- | --- | --- | --- | --- | --- | --- | --- | --- | --- |
|  |  |  |  |  |  |  | Before | After | Kept | Lost | Newly acquired | Before | After |
|  |  |  |  |  |  | 1 | 319 | 715 | 114 | 205 | 601 | 4.4 | 5.1 |
|  |  |  |  |  |  | 2 | 370 | 848 | 131 | 239 | 717 | 3.7 | 5.0 |
|  |  |  |  |  |  | 3 | 237 | 1179 | 93 | 144 | 1086 | 3.9 | 6.2 |
|  |  |  |  |  |  | 4 | NA | 841 | NA | NA | NA | NA | 5.5 |
|  |  |  |  |  |  | 5 | 314 | 843 | 92 | 222 | 751 | 3.1 | 5.2 |
|  |  |  |  |  |  | 6 | NA | 1381 | NA | NA | NA | NA | 6.5 |
|  |  |  |  |  |  | 7 | NA | 739 | NA | NA | NA | NA | 5.4 |
|  |  |  |  |  |  | 8 | NA | 3574 | NA | NA | NA | NA | 8.5 |
| Touched surface |  |  |  |  |  | mean | 310.0 | 1265.0 | 107.5 | 202.5 | 788.8 | 3.7 | 5.9 |
|  |  |  |  |  |  | sd | 54.9 | 960.9 | 18.7 | 41.4 | 208.3 | 0.5 | 1.2 |

Subway passenger microbiome profiles converged after traveling. The number of taxa shared among the eight passengers increased after traveling without handwashing (Figs. 5b, 6, S4b). A constrained analysis of principal coordinates (CAP) supported the microbial convergence after a subway ride, showing that the subject identifier variable (SubjectID) followed by the travel variable (before and after) significantly explained group segregation (*p* = 0.001, F = 2.4 and *p* = 0.006, F = 2.0, respectively; ANOVA-like permutation test for CAP, Fig. 5c). The handwashing procedure analysis showed similar results (*p* = 0.003, F = 1.9), although the SubjectID variable reduced to marginal significance (*p* = 0.049, F = 1.3; Fig. S4c). The convergence of the subject microbial composition after traveling can also be visualized in an NMDS ordination; SubjectIDs after traveling were closer to each other (Fig. 6). Additionally, they became closer to the subway surface profiles, suggesting higher similarities with the surface microbiome, particularly evident for the hand-washing group.

**Figure 6.**
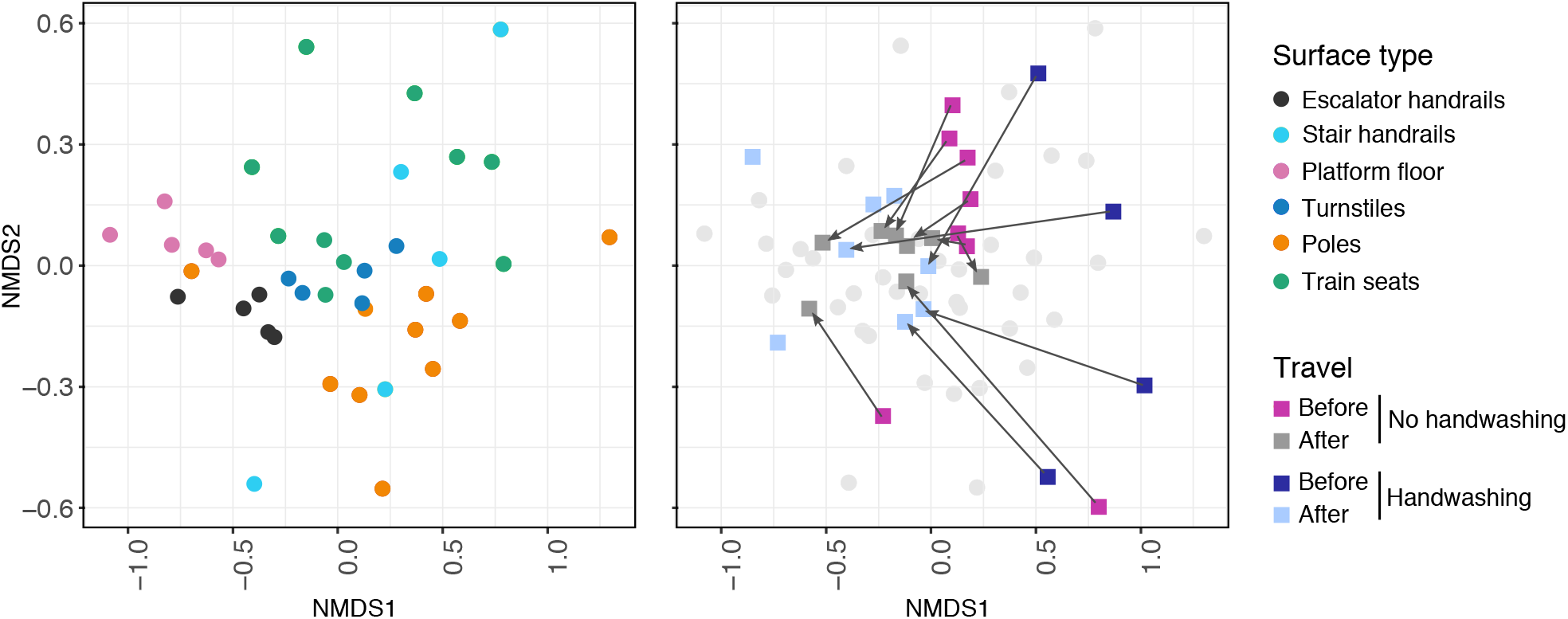
Hand microbiome composition converges after traveling. Non-multidimensional scaling (NMDS) ordination bi-plot performed with Bray Curtis distance for no handwashing and handwashing and for different surface types, at the genus level. Arrows connect the same subjects from before to after traveling. Unpaired dots are samples without matching before or after travel comparison because of low metagenomic DNA yield (washed hands).

**Figure S4.**
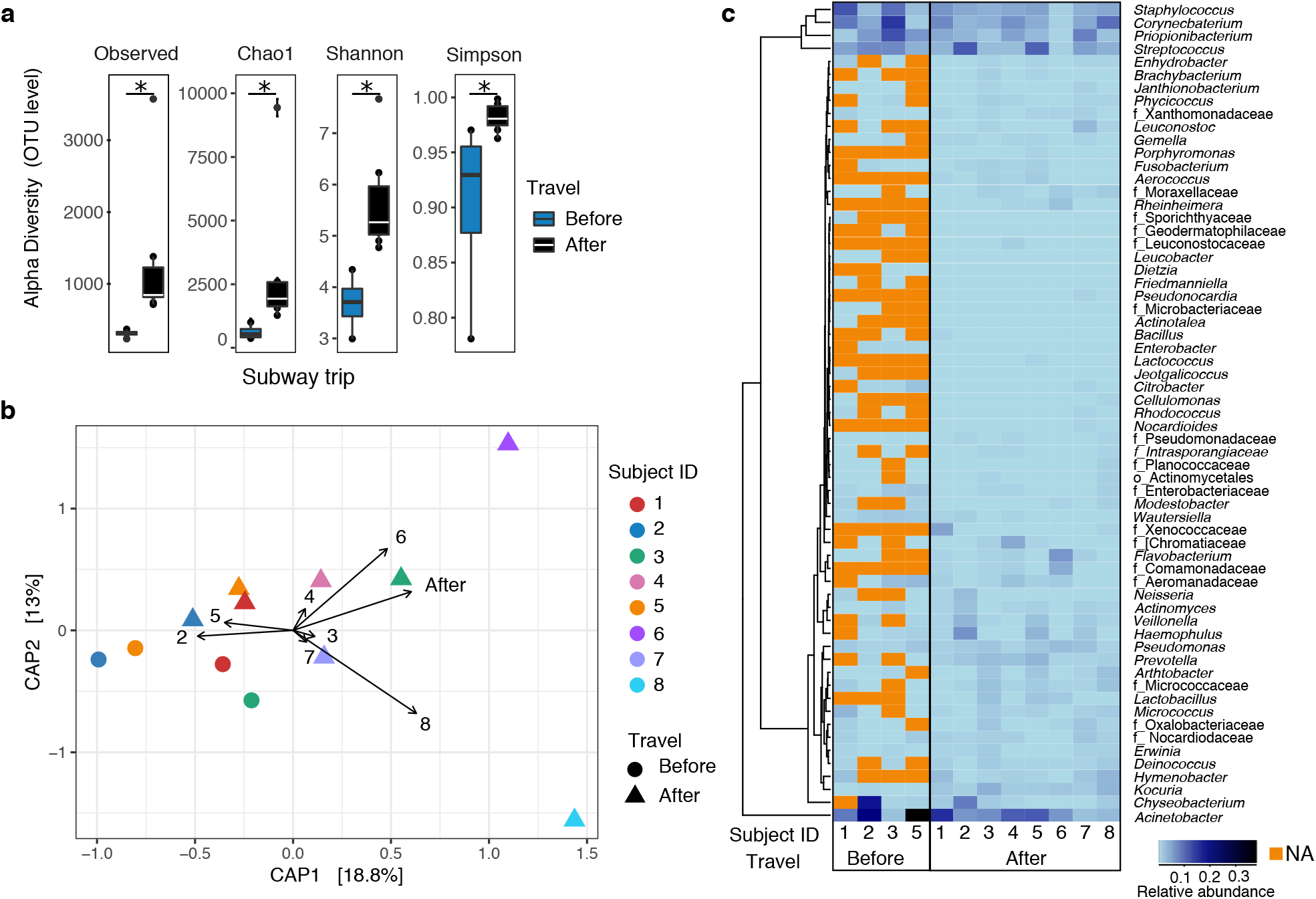
Changes in the passenger hands microbiome before and after traveling with handwashing. (a) Alpha diversity is increased (* *p* < 0.020). (b) Heatmap showing the relative abundance of all common taxa at the genus level, before or after traveling per SubjectID, denoted by a number. Row dendrogram arrangement based in Manhattan distance. (c) Constrained analysis of principal coordinates (CAP) at the genus level, showing significant segregation of SubjectID and Travel variables (*p* = 0.049, F = 1.3 and *p* = 0.003, respectively, F = 1.9, ANOVA-like permutation test for CAP).

**Figure S5.**
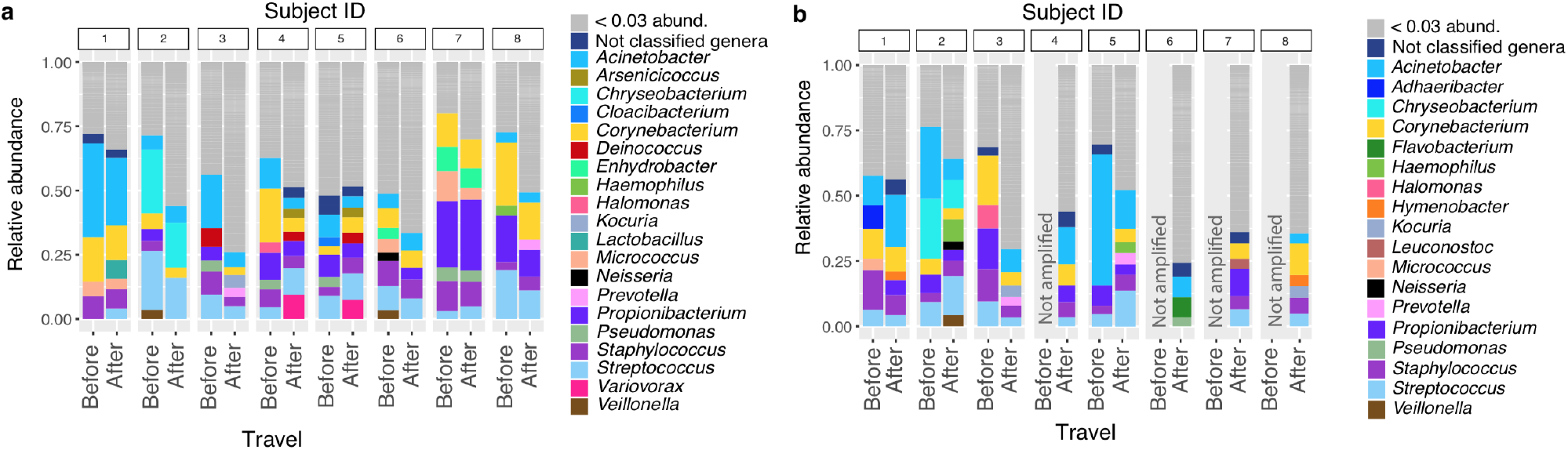
Taxa summary at the genus level per SubjectID. Each subject fingerprint is preserved after traveling (a) Before and after traveling without handwashing. (b) With handwashing. Missing bars come from samples not sequenced due to low DNA biomass.

In summary, subway traveling increased hand microbial diversity and promoted passenger microbiome convergence. Although handwashing before traveling had an immediate effect on microbiome profiles, diversity and composition reached similar characteristics than after traveling without handwashing.

### Pathogenic bacteria

We did not detect fecal indicators such as *Escherichia coli* in any surface or hand samples. However, other coliform genera were present: *Klebsiella* (13% prevalence in all samples), *Enterobacter* (90% prevalence), and *Citrobacter* (90% prevalence), with no differences among surface types or traveling variables.

A total of 59 (mean of 19±6 per sample) potentially pathogenic bacteria were identified (we used the non-rarified data set) based on a list of 554 (543 genera + species and 11 genera) (Table S9). We used two sources to identify pathogens: The Taylor, Latham, & Woolhouse list (Taylor et al., 2001) and from the National Institute of Allergy and Infectious Diseases (NIAID) Emerging Infectious Diseases/Pathogens list {NIAID Emerging Infectious Diseases, 2014). The number of potentially pathogenic taxa differed among surface types (p = 0.021, Kruskal-Wallis), being higher in escalator handrails and lower in seats (29 vs. 15, Table S10). Passenger hands before traveling had a mean of 15 potentially pathogenic taxa, which was increased after traveling (*p* < 0.02, Wilcoxon test, Table S11). Notably, after traveling, passengers equaled the count of potentially pathogenic taxa to the subway surface mean count (21 taxa). Although the increase in microbial diversity increases the probability of contact with pathogenic bacteria, the volunteers were healthy and had carried the pathogenic taxa before entering the subway.

**Table S10.** Potentially pathogenic taxa detected on subway surfaces.

| Surface sample ID | Number of pathogenic taxa (%)<sup>*</sup> |  |  |  |  |  |
| --- | --- | --- | --- | --- | --- | --- |
|  | Seats | Escalator handrails | Turnstiles | Poles | Platform floor | Regular stair handrails |
| 1 | 18 | 30 | 29 | 16 | 24 | 13 |
| 2 | 19 | 26 | 14 | 22 | 24 | 15 |
| 3 | 21 | 31 | 15 | 24 | 23 | 28 |
| 4 | 10 | 31 | 20 | 18 | 19 | 9 |
| 5 | 8 | 26 | 28 | 15 | 19 | 26 |
| **mean** | **15.2** | **28.8** | **21.2** | **19.0** | **21.8** | **18.2** |
| **p value<sup>\*\*\*</sup>** | **0.021** |  |  |  |  |  |

<sup>\*</sup>554 Genus+Species from two list: Taylor's list (Taylor et al, 2001) and National Institute of Allergy and Infectious Diseases (NIAID) Emerging Infectious Diseases/Pathogens list
<sup>\*\*</sup> Kruskal-Wallis test

**Table S11.** Potentially pathogenic taxa before and after subway traveling. Two conditions were evaluated: entering the subway without washed hands and with washed hands.

| Subject ID | Without handwashing |  | Handwashing |  |
| --- | --- | --- | --- | --- |
|  | Number of potentially pathogenic taxa<sup>*</sup> |  | Number of potentially pathogenic taxa<sup>*</sup> |  |
|  | Before | After | Before | After |
| 1 | 16 | 17 | 6 | 15 |
| 2 | 23 | 24 | 12 | 21 |
| 3 | 16 | 29 | 8 | 22 |
| 4 | 14 | 23 | NA | 19 |
| 5 | 12 | 21 | 15 | 21 |
| 6 | 17 | 18 | NA | 20 |
| 7 | 16 | 18 | NA | 13 |
| 8 | 9 | 25 | NA | 30 |
| **Mean or proportion** | **15.4** | **21.9** | **10.3** | **20.1** |
| **p value<sup>\*\*</sup>** | **0.005** |  | **0.017** |  |

<sup>\*</sup>554 Genus+Species from two list: Taylor's list (Taylor et al, 2001) and National Institute of Allergy and Infectious Diseases (NIAID) Emerging Infectious Diseases/Pathogens list
<sup>\*\*</sup> Wilcoxon test

### ASVs analysis

We reanalyzed the central questions of this study using amplicon sequence variants (ASV). Both, ASV and OTU are methodologies for data reduction. While, ASVs are generated by parsing identical sequences and elimination of rare sequences, OTUs are generated by sequence clustering. Presenting ASV approach may allow a more complete understanding of the data since it can provide strain resolution. Microbial diversity at the ASV level showed similar results than for the OTU analysis in alpha and beta diversity for subway surfaces and hands before and after traveling. Previous subway studies have reported the presence of *E. coli,* or even *Escherichia,* as opposed to this study. The ASV analysis detected 13 *Escherichia-Shigella* in some surface and passenger samples without any particular pattern. We compared regular and women-only wagons, identifying the presence and changes in the relative abundance of specific vaginal-associated bacteria based on the Silva database. This database includes a higher number of vaginal-associated bacteria than GreenGenes. Similar to the OTU analysis, no differences were found (Table S12, Fig. S6, Table S13).

**Table S12.**
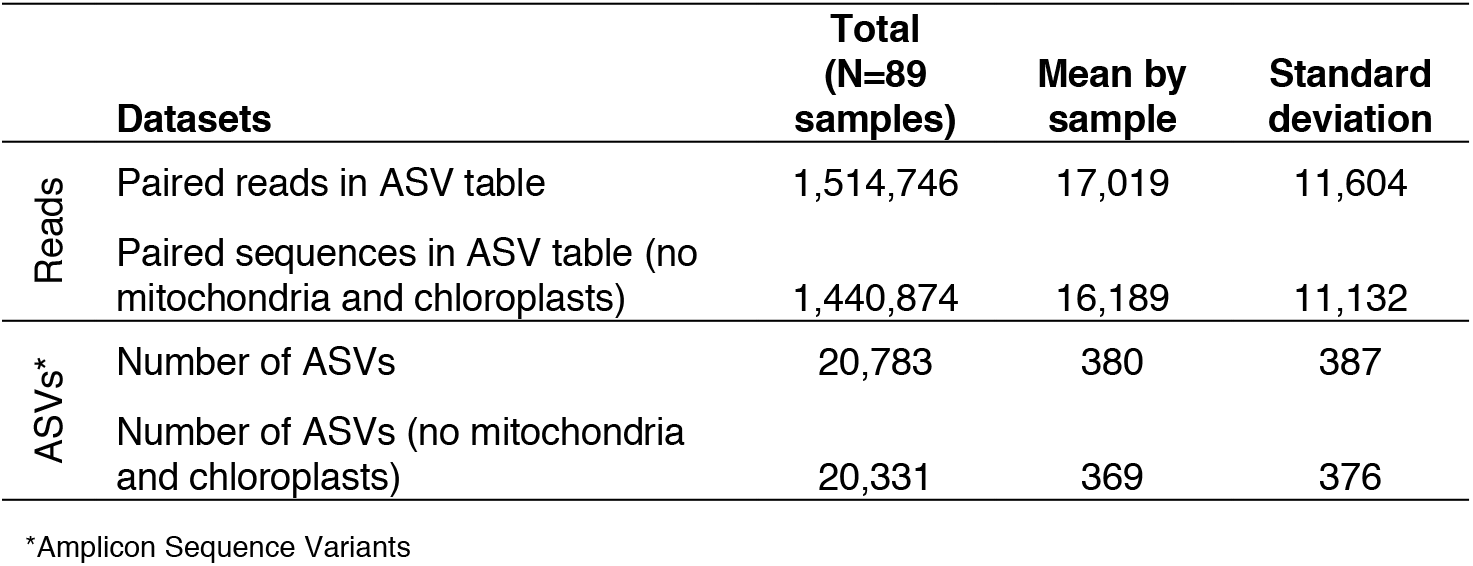
Summary of sequences and ASVs in 89 samples

**Figure S6.**
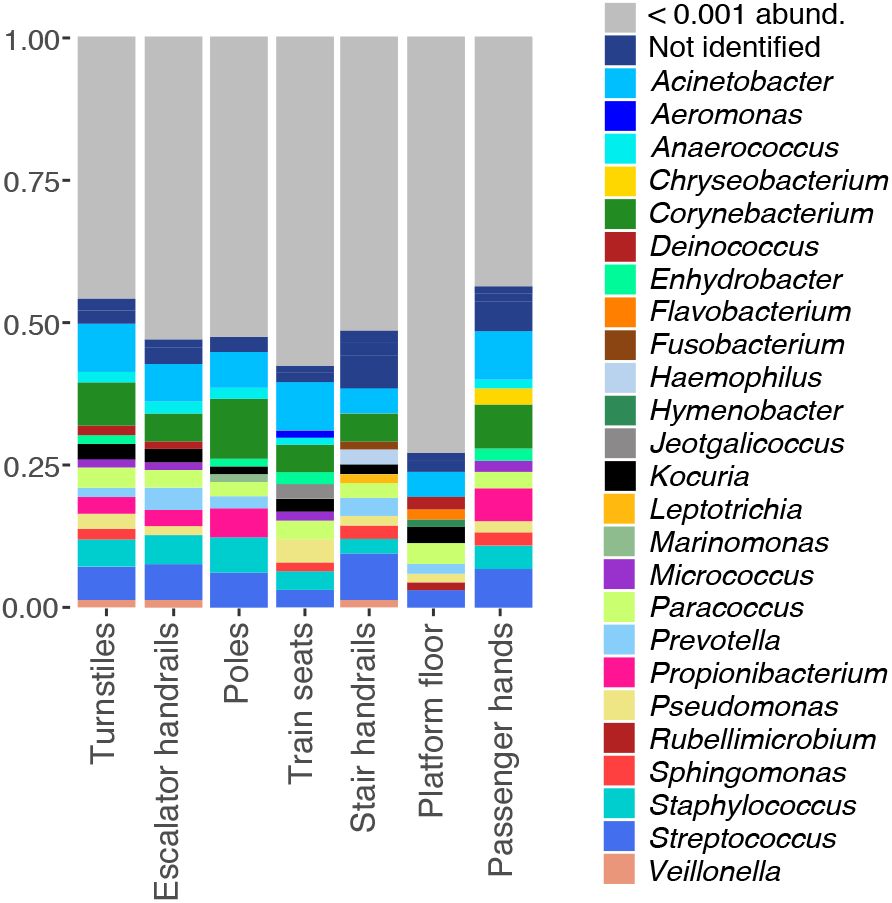
Taxa summary at genus level from ASVs generated with DADA2 and taxonomy assign based on Silva Database. Similar to the OTU analysis, the most abundant phyla were Proteobacteria (37%), Firmicutes (24%), Actinobacteria (20%), and Bacteroidetes (9.3%). However, Cyanobacteria (3.3%, no chloroplast) appeared in the fifth position. The most abundant ASVs were *Acinetobacter lwoffii* (0.72%), *Streptococcus* sp. (0.68%), *Streptococcus* sp. (0.59%), *Acinetobacter lwoffii* (0.56%), and *Propionibacterium acnes* (0.56%). A total of 17 ASVs named as archaea were identified *(Methanobrevibacter, Candidatus Nitrososphaera SCA1170, Candidatus Nitrososphaera SCA1145, Methanosaeta vadinCA11, Methanosaeta, Natronococcus, Methanobacterium, Halococcus*, among other not identified genera).

**Table S13.** Number of vaginal-associated taxa as a female environmental indicator.

| Taxa (ASV)* | Poles |  | Train seats |  |
| --- | --- | --- | --- | --- |
|  | Regular (N=5) | Women-only (N=5) | Regular (N=5) | Women-only (N=5) |
| *Lactobacillus crispatus* | 0 | 0 | 0 | 0 |
| *Lactobacillus iners* AB-1 | 1 | 2 | 1 | 1 |
| *Lactobacillus gasseri* | 0 | 0 | 1 | 0 |
| *Lactobacillus jensenii* | 0 | 0 | 0 | 0 |
| *Atopobium vaginae* | 0 | 1 | 0 | 0 |
| *Sneathia sanguinegens* | 0 | 0 | 0 | 0 |
| *Sneathia amnii* | 0 | 0 | 0 | 0 |
| *Prevotella bivia* | 1 | 1 | 0 | 1 |
| *Prevotella disiens* | 0 | 0 | 0 | 1 |
| **Sum** | **2** | **4** | **2** | **3** |

\* Amplicon sequence variants taxonomic assignment was performed using the Silva database.

## Discussion

Microbial subway surfaces have been characterized previously in New York, Boston, Oslo, and México City (Afshinnekoo et al., 2015; Hsu et al., 2016; Gohli et al., 2019; Hernández et al., 2019). Additionally, the hand microbiome has been explored for Hong Kong subway passengers (Kang et al., 2018). However, no study has integrated subway surfaces and their taxonomic contribution to the passengers’ hands.

The Mexico City subway showed different microbial compositions among surface types. Differences in the material physical composition and the type of human body interaction may be shaping these profiles (Hsu et al., 2016; Hernández et al., 2019). Although escalator handrails and poles are surfaces typically wrapped by the passengers’ hands, we found substantial differences in diversity and compositions among them. The higher porosity in the escalator rubber grips may provide a higher contact surface and a higher faculty of harboring nutrient particles that facilitate bacterial growth. Additionally, it may serve as a humidity reservoir for bacteria (Verdier, et al., 2014). On the contrary, poles are polished metal surfaces that reduce bacterial adherence and persistence. Differences in the type of usage is also a fundamental microbiome-shaping factor. The platform floor, with the most distinctive composition, receives soil and dust particles carried in via shoes, while seats are impacted by the commuters’ clothes. Turnstiles, handrails, and poles show the microbial input of hands and clothes.

We also explored whether regular and women-only wagons displayed differentiated microbiomes, based on the previously reported sex-based microbial differences and building-occupiers microbial associations (Fierer et al., 2008; Ross et al., 2017; Takagi et al., 2019; Song et al., 2013; Lax et al., 2014). We did not find distinctions between regular and women-only wagons. Microbiome sex-based signals in trains may be hidden by the high intrinsic diversities of the subway surfaces. A female signal would probably require contact with a low, exposed body part *(e.g.,* thighs or urogenital area) (Ross et al., 2017; Gibbons et al., 2015; Flores et al., 2011). Additionally, these wagons are intermittently occupied by women. The first wagons are for women only, so each time each train reaches a terminal station, there is a railroad switch, reversing train wagon order. A 20-30-min ride would probably be not long enough to build a female microbial fingerprint. Additionally, a possible PCR primer bias might be limiting the detection of vaginal species (*e.g., Lactobacillus* spp.), which may be difficult accessing this comparison.

### The passenger microbial input

We described the poles as the least bacterial diverse surface but as the most frequent passenger wrap surface (97%). Such a highly perturbed surface might be of particular interest in controlling microbial dispersal with health implications. In this study, we observed that pole cleaning effectively removed microbes. However, we observed microbial resettling with few passenger interactions with bacterial richness and UFC counts similar to pre-cleaning levels. The fact that microbes rapidly resettle suggests that cleaning is not effective over time. Although the smoothness of the pole may avoid microbial accumulation, it might promote a high microbial exchange rate. In practical terms, the passenger microbes are rapidly wiped out by the next passenger.

We observed changes in the pole microbial compositions across time groups and did not detect an evident sign of ecological succession. However, we cannot rule out a slow succession process. Gibbons et al. (2015) have followed the microbial colonization on restroom floors and observed an early successional community composition within 8 h and a late-successional state over weeks to months. The observed high perturbation frequency of poles and the scarce deposition area of its material may impair the development of a community structure over time (Connell and Slatyer, 1977).

Further studies of cleaning porous surfaces such as escalator handrails (used by 86% of the passengers) may be essential to prevent microbial spreads; however, microbial removal efficiency must be tested. We also propose to pay particular attention to the cleaning of the floor, as floor surfaces are highly diverse and the dust can be easily lifted with air and breathed in by passengers. Alternatively, the use of self-cleaning building materials may be a long-term strategy for controlling bacterial colonization on surfaces (Benedix et al., 2000).

### Passenger hand microbiome after a subway ride

We showed that the hands are the primary means of interaction with subway surfaces. Additionally, we observed that the face/head area was the area most commonly touched by users (73%). From this area, the mucosa has a particular health relevance. Interaction with mucosa was relatively high in prevalence (17%, 10-min trip). In contrast to the skin, with a robust mechanical barrier, the mucosa is an exposed area for external agents which are directly in contact with the immune system. Although recognition and protection against environmental agents are continually occurring in this area, the mucosa can also be vulnerable to disruption and microbial colonization. A persistent establishment would imply interactions with the immune system (Edmonds-Wilson et al, 2015), while a transient establishment involves bacterial dispersal to other surfaces.

Bacterial adherence ability plays a vital role in the microbial exchange. It might be influenced by temperature and pressure conditions as well as by the hydrophobic-hydrophilic properties of the interacting surfaces (Liu et al., 2004). These properties may vary by surface characteristics and passenger hand microenvironment (pH, moisture, sebum level). Based on our data, touching escalator handrails is equivalent to shaking the hands of three to four different people.

After a subway trip, the passenger hand microbiome increased in diversity. This increment is consistent with the higher diversity found on subway surfaces, which may be built by the contribution of a high passenger influx and soil presence (Hernández et al., 2019). The hand from one person is a heterogeneous microbial source with high inter-individual variation (Fierer et al., 2008). We also detected commuter variation; only 7% of the genera were shared among passenger hands before traveling. This high inter-variation may be due to intrinsic factors such as age (Flores et al., 2014; Song et al., 2013), sex (Takagi et al., 2019), and extrinsic lifestyle-dependent factors: use of skincare products, pet ownership, allergies, alcohol consumption, time spent outdoors (Ross et al., 2017).

We showed that hand microbial composition converged among passengers after traveling. This convergence means that one trip is enough to perceive the effect of building cohabitation (Lax et al., 2014; Ross et al., 2017; Song et al., 2013). Changes in the hand microbiome were expected, as hands were constantly interacting with different surfaces. Hands show higher temporal variability than other body sites (reviewed in Edmonds-Wilson et al., 2015).

Besides diversity changes due to travel, we observed microbial fingerprint preservation within passengers. Highly abundant taxa persist within subjects (Caporaso et al., 2011), while transient bacteria are the main variability source (Flores et al., 2014). This has also been observed in volunteers that were sampled and resampled at 4 to 6 months later, with no significant change over time (Grice et al., 2009).

### Potential pathogens and the safeness of riding the subway

We identified the presence of potentially pathogenic taxa. After traveling, the number of potentially pathogenic species increased, reaching the same levels as the subway surfaces. Potential pathogens include organisms that do not usually cause disease in a healthy population but that are relevant to people with open wounds, immunocompromised, or the elderly because of their intrinsic decline of immune responsiveness (Wick & Grubeck-Loebenstein, 1997). Potentially pathogenic taxa have also been reported in other surfaces such as ATMs (Bik et al., 2016) or kitchen sponges (Marotta et al., 2018). In our study, the identification of such species *(e.g., S. aureus, C. perfringens)* is not surprising (see also Cave et al., 2019). These species can be found in healthy humans; for example, *S. aureus* colonizes 70% of the healthy Mexican population (Hamdan-Partida et al., 2010), and *C. perfringens* is part of the healthy intestinal human microbiota (Palmer et al., 2007). However, some strains are highly pathogenic or have antibiotic resistance (*e.g.,* methicillin-resistant *S. aureus*), which invites to further explore viability and strain pathogenic ability with targeted studies.

## Conclusions

We detected the effect of building cohabitation on passengers of the Mexico City subway. Each time a passenger travels in the subway, she or he leaves some bacteria and brings others. Passengers become similar to the subway surfaces, and they are more alike among each other after traveling. Each passenger’s microbial fingerprint is preserved, mostly explained by high-abundance taxa. Although most bacteria will not persist, traveling in the subway is a way of sharing our microbes.

Poles are the most touched surface in the Mexico City subway. Pole cleaning reduces microbial richness and diversity. However, the amount of UFC is quickly restored within 5 min after pole cleaning. Even so, the microbial composition is not resettled within 48 hours. We think that the lack of restoration of the initial microbial community is due to the variability of the hosts’ microbiomes and the lack of persistence of rare taxa.

## Methods

### Sampling

Sampling was performed in autumn 2018. Samples were taken from turnstiles, stair handrails, escalator handrails, platform floor, train poles, and seats. The latter two were sampled from two different train wagons: the regular wagon (used by men and women) and the women-only wagon (used by women, disabled people, and the elderly). A total of 97 samples were collected, and 89 were successfully processed (Table S1). Sampling was performed by swabbing each surface of around 100 cm^2^ for 20 seconds with a pre-moistened nylon-flocked swab (COPAN FLOQSwabs™), and samples were preserved in transport media (Tris 20 mM, EDTA 10 mM pH 7.5). Samples were kept in ice for less than 12 h until freezing at −80°C. Line and station names, time, temperature, and relative humidity were registered.

The impact of a subway trip on the passengers’ microbiomes was determined by swabbing the right hand of eight informed volunteers. Volunteers were sampled before and after traveling at a regular weekday. Subjects arrived at the starting point in the morning, not having used the subway as a transportation means. The subway trip included traveling 11 stations across three different subway lines, including two-line transferences. It was a circular route, so they would arrive at the starting point. Each volunteer was indicated to get in contact with particular surfaces (at least twice) and to avoid touching others. Surfaces touched by each volunteer are summarized in Tables S6 and S7. Hands were sampled after completing the trip. Additionally, volunteers were asked to wash their hands for a period of 30 s with liquid soap (DIAL^®^ neutral) and distilled water. Immediately after this, hands were sampled and then resampled at the end of the trip.

### Surfaces cleaning

To describe the microbiome colonization after surface decontamination, five poles from the same wagon were chosen (mixed wagon) in the morning of a regular weekday. Areas to sample were defined with a template divided into five areas of 100 cm^2^ each. Samples were taken before and after cleaning (pre- and post-cleaning). A Lysol wipe was used to energetically scrub the surface, and then, a wet sterile gauze was used to remove the cleaning product excess. Post-cleaning samples were swabbed immediately after the cleaning of each surface. The remaining samples were taken longitudinally at five time-points (0, 0.5, 2, 8, and 48 h). We did not sample twice in the same area to avoid affecting the microbiome composition of the following time points.

Surface cleaning was also analyzed by CFU counting. For this, 10-12 poles were sampled as described above, in seven post-cleaning time points: 0 h, 5 min, 10 min, 20 min, 0.5 h, 1 h, and 2 h. Two positive controls were also swabbed, pre-cleaning samples and 2 h without cleaning. Bacteria were incubated in LB-agar medium for 36 h at 30°C.

### Observational patterns during a subway trip

A total of 120 passengers (67 adults and 53 elderly) were randomly picked and observed from the start to the end of each trip. All interactions with their environment were documented during traveling. Additionally, observations were also made in the stations. The number of passengers touching the handrails of escalators and stairs were documented in 5 and 11 different stations, respectively. Differences between stairs going up or down were also explored, while the chosen escalators were only going up.

### DNA extraction

Metagenomic DNA was extracted using a MoBio PowerSoil Kit (MoBio Laboratories, Solana Beach, CA) with small modifications. Half of the beat material from each tube was poured in a new one. Volumes were adjusted to preserve proportions of solutions/samples. In total, 125 μL of the sample were used, 30 μL of C1 solution and 50 μL of phenol: chloroform 1:1 were mixed in the beat tube. Further steps were performed according to the instructions of the MoBio PowerSoil Kit.

Region V3-V4 from the 16S rRNA gene was amplified using primers 341F (CCTACGGGNGGCWGCAG) and 805R (GACTACHVGGGTATCTAATCC). Libraries were obtained following the MiSeq™ Illumina^®^ protocol. Samples without PCR amplification were discarded. All samples corresponded to the ones taken just after cleaning the surfaces. The PCR was performed in triplicates, using 0.15 ul of Phusion DNA polymerase™, 3 μL Buffer 5x, 2.5 μL dNTP (3 mM), 1 μL forward primer (5 pmol/μL), 1 μL reverse primer (5 pmol/μL), 1-4 μL DNA, and water up to the final volume of 15 ul per reaction. The PCR reaction was initiated at 98°C, 30 s, followed by 35 cycles of 92°C for 10 s, 53°C for 30 s, 72°C for 40 s, and a final extension step at 72°C for 5 min. Blank samples were used as negative controls. The three PCR reactions per sample were combined and purified using the High Pure PCR Product Purification Kit of ROCHE™ (Roche Diagnostics GmbH, Mannheim, Germany). Sequencing was performed using the MiSeq™ Illumina^®^ (2 x 300 bp) platform at the Laboratory of Genomic Services from the National Laboratory of Genomics for Biodiversity in Irapuato, México. The DNA concentration was measured with a NanoDrop microvolume spectrophotometer.

### Bioinformatics processing

Amplified reads were pair-ended using the Context-Aware Scheme for Paired-End Read (CASPER) (Kwon, Lee, & Yoon, 2014). Sequences were clustered at 97% of identity using cd-hit-est (Li & Godzik, 2006), and pick_rep_set.py from QIIME (v. 1.9) (Caporaso et al., 2010) was used to pick representative sequences from each cluster. Chimera sequences and singletons were removed. The taxonomy assignment was done with QIIME (v. 1.9) (Caporaso et al., 2010) using parallel BLAST (Camacho et al., 2009) and the GreenGenes database (DeSantis et al., 2006). Finally, chloroplasts and mitochondria were filtered from the OTU table. These steps were processed with default parameters, and the analyzed samples were rarified at 6,242 sequences per sample.

Additionally, amplicon sequence variants (ASVs) were generated using the DADA2 algorithm (Callahan et al., 2016). Reads 341F-804R were extracted from the 16S Silva database (Quast et al., 2013) to train a Naive Bayes classifier (Wang et al., 2007).

### Pathogenic taxa

The presence of potentially pathogenic taxa was analyzed comparing the obtained bacteria with a list comprising 554 taxa (genus or genus + species) from two sources, from a previous study (Taylor et al., 2001) including bacteria related to adverse health outcomes and from the National Institute of Allergy and Infectious Diseases (NIAID, 2014) Emerging Infectious Diseases/Pathogens list (Green et al., 2007).

### Data analysis

Data analysis and plot generation were performed using *phyloseq* (McMurdie & Holmes, 2013) and *ggplot2* (Wickham, 2009) in R version 3.5.1 (R Development Core Team, 2011). Beta diversity was visualized with canonical analysis of principal coordinates (CAP) and non-metric dimensional scaling (NMDS) with Bray Curtis dissimilarities. Comparison among groups was performed with the vegan package (Oksanen et al., 2013) from R, using the adonis function, which performs a permutational multivariate analysis of variance (PERMANOVA) using distance matrices with 999 permutations. Group dispersion was examined with multivariate homogeneity of group dispersions with the betadisp R function. PERMANOVA pairwise comparison was performed with the pairwise.adonis function (Martinez Arbizu, 2017) in *devtools* package with default parameters and adjusted p values with the false discovery rate (FDR) method. Non-parametric comparisons were performed with the Kruskal-Wallis test and pairwise comparison with Dunn’s Test of Multiple Comparisons Using Rank Sums, dunn.test function from the R base library. Non-parametric two-group comparison was performed with the Wilcoxon test.

## Declarations

### Consent for publication

No personal identifiers were included in the available data. Sampling permits were granted by the metro “User Support Manager” (Gerencia de Atención al Usuario, del Sistema de Transporte Colectivo).

### Availability of data and materials

The datasets supporting the conclusions of this article are available in the Sequence Read Archive (SRA) at the NCBI, with the submission number SUB6707685 (https://submit.ncbi.nlm.nih.gov/subs/sra).

### Ethical approval

Comisión de Ética Académica y Responsabilidad Científica de la Facultad de Ciencias, UNAM (CEARC/Bioética/09042020).

### Authors’ contributions

DVR, CG-C, AMH, and MP designed the study. DV-R, CG-C, and AMH collected the data. DV-R processed and analyzed the data. LDA and MP conceived the study. DV-R, LDA, and MP wrote the paper.

## Acknowledgments

We thank the subway customer service (*Sistema de Transporte Colectivo, Cd. Mx.*) for providing sampling permissions. We also thank the participation of graduate students, for providing their time, effort, and letting us swab their hands. We thank Fabricio Martinez and Carla Cicero from the Autonomous University of Mexico City (UACM) for their support in the sample collection.

---

**Table S4:** Relative humidity, temperature, collection date, DNA concentration, and number of raw sequences per sample.

| sample_ID | sample_type | rel_air_humidity | air_temp | collection_date | DNA_UV<br>(ng_uL) | raw_assembled_<br>sequences |
| --- | --- | --- | --- | --- | --- | --- |
| LA5 | skin_right_hand | 44 | 19 | 22-Nov-2018 | 5.08 | 33893 |
| LD5 | skin_right_hand | 38 | 22 | 22-Nov-2018 | 14.74 | 60172 |
| SD5 | skin_right_hand | 45 | 19 | 22-Nov-2018 | 86.29 | 40185 |
| SA5 | skin_right_hand | 40 | 22 | 22-Nov-2018 | 68.39 | 38808 |
| LA2 | skin_right_hand | 44 | 19 | 22-Nov-2018 | 1.27 | 77430 |
| LD2 | skin_right_hand | 38 | 22 | 22-Nov-2018 | 4.84 | 131270 |
| SA2 | skin_right_hand | 40 | 22 | 22-Nov-2018 | 8.74 | 48905 |
| SD2 | skin_right_hand | 45 | 19 | 22-Nov-2018 | 15.27 | 54193 |
| LD3 | skin_right_hand | 38 | 22 | 22-Nov-2018 | 19.09 | 198903 |
| SD3 | skin_right_hand | 45 | 19 | 22-Nov-2018 | 21.56 | 54015 |
| LA3 | skin_right_hand | 44 | 19 | 22-Nov-2018 | 31.57 | 91923 |
| SA3 | skin_right_hand | 40 | 22 | 22-Nov-2018 | 1.55 | 33912 |
| LD1 | skin_right_hand | 38 | 22 | 22-Nov-2018 | 2.09 | 87439 |
| SA1 | skin_right_hand | 40 | 22 | 22-Nov-2018 | 4.78 | 44297 |
| LA1 | skin_right_hand | 44 | 19 | 22-Nov-2018 | 1.07 | 15136 |
| SD1 | skin_right_hand | 45 | 19 | 22-Nov-2018 | 126.42 | 34589 |
| LD6 | skin_right_hand | 38 | 22 | 22-Nov-2018 | 23.57 | 77792 |
| SD6 | skin_right_hand | 45 | 19 | 22-Nov-2018 | 1.57 | 42246 |
| SA6 | skin_right_hand | 40 | 22 | 22-Nov-2018 | 10.39 | 33103 |
| LD4 | skin_right_hand | 38 | 22 | 22-Nov-2018 | 74.66 | 86562 |
| SD4 | skin_right_hand | 45 | 19 | 22-Nov-2018 | 22.07 | 54146 |
| SA4 | skin_right_hand | 40 | 22 | 22-Nov-2018 | 45.90 | 34314 |
| LD7 | skin_right_hand | 38 | 22 | 22-Nov-2018 | 112.31 | 44363 |
| SA7 | skin_right_hand | 40 | 22 | 22-Nov-2018 | 2.97 | 30958 |
| SD7 | skin_right_hand | 45 | 19 | 22-Nov-2018 | 28.73 | 36741 |
| LD8 | skin_right_hand | 38 | 22 | 22-Nov-2018 | 206.29 | 101256 |
| SA8 | skin_right_hand | 40 | 22 | 22-Nov-2018 | 4.40 | 19890 |
| SD8 | skin_right_hand | 45 | 19 | 22-Nov-2018 | 3.77 | 44467 |
| TP1 | turnstile | 47 | 22 | 29-Oct-2018 | 55.84 | 43941 |
| TP2 | turnstile | 48 | 21 | 29-Oct-2018 | 26.90 | 23529 |
| TP3 | turnstile | 49 | 21 | 29-Oct-2018 | 68.89 | 39309 |
| TP4 | turnstile | 52 | 20 | 29-Oct-2018 | 38.60 | 39106 |
| TP5 | turnstile | 52 | 20 | 29-Oct-2018 | 5.43 | 54493 |
| PR3 | stairs_handrail | 31 | 26 | 20-Nov-2018 | 50.89 | 121669 |
| PR1 | stairs_handrail | 34 | 24 | 20-Nov-2018 | 34.96 | 35905 |
| PR5 | stairs_handrail | 31 | 28 | 20-Nov-2018 | 98.64 | 53191 |
| PR4 | stairs_handrail | 36 | 27 | 20-Nov-2018 | 1.49 | 62671 |
| PR2 | stairs_handrail | 38 | 24 | 20-Nov-2018 | 2.44 | 48445 |
| PM3 | scalator_handrail | 33 | 26 | 20-Nov-2018 | 11.35 | 82668 |
| PM1 | scalator_handrail | 34 | 24 | 20-Nov-2018 | 18.41 | 71606 |
| PM5 | scalator_handrail | 31 | 28 | 20-Nov-2018 | 13.06 | 64366 |
| PM4 | scalator_handrail | 36 | 27 | 20-Nov-2018 | 28.04 | 79962 |
| PM2 | scalator_handrail | 38 | 24 | 20-Nov-2018 | 33.05 | 87440 |
| BF1 | pole | 44 | 27 | 31-Oct-2018 | 1.80 | 49284 |
| BF2 | pole | 43 | 27 | 31-Oct-2018 | 1.80 | 24771 |
| BF3 | pole | 44 | 27 | 31-Oct-2018 | 24.17 | 41976 |
| BF5 | pole | 46 | 27 | 31-Oct-2018 | 1.57 | 50129 |
| B22 | pole | 49 | 28 | 29-Oct-2018 | 59.81 | 38287 |
| B23 | pole | 39 | 32 | 29-Oct-2018 | 86.71 | 41361 |
| B24 | pole | 37 | 32 | 29-Oct-2018 | 1.72 | 33335 |
| B25 | pole | 37 | 32 | 29-Oct-2018 | 20.59 | 76069 |
| Bc1 | pole | 56 | 20 | 29-Oct-2018 | 2.36 | 51959 |
| Bc2 | pole | 46 | 28 | 29-Oct-2018 | 7.40 | 24060 |
| Bc4 | pole | 46 | 29 | 29-Oct-2018 | 2.02 | 28946 |
| B41 | pole | 46 | 26 | 31-Oct-2018 | 1.83 | 45469 |
| B42 | pole | 46 | 26 | 31-Oct-2018 | 1.74 | 42237 |
| B45 | pole | 46 | 27 | 31-Oct-2018 | 1.14 | 72208 |
| B81 | pole | 58 | 25 | 29-Oct-2018 | 7.22 | 92576 |
| B82 | pole | 48 | 26 | 29-Oct-2018 | 3.15 | 41792 |
| B83 | pole | 48 | 27 | 29-Oct-2018 | 1.17 | 28386 |
| B84 | pole | 55 | 27 | 29-Oct-2018 | 1.87 | 36519 |
| B85 | pole | 47 | 29 | 29-Oct-2018 | 2.33 | 118089 |
| BP1 | pole | 56 | 20 | 29-Oct-2018 | 39.34 | 101097 |
| BP2 | pole | 46 | 28 | 29-Oct-2018 | 14.00 | 72347 |
| BP3 | pole | 50 | 29 | 29-Oct-2018 | 38.87 | 244572 |
| BP4 | pole | 46 | 29 | 29-Oct-2018 | 5.30 | 86486 |
| BF4 | pole | 47 | 27 | 31-Oct-2018 | 65.70 | 24143 |
| B21 | pole | 40 | 29 | 29-Oct-2018 | 21.78 | 18315 |
| Bc3 | pole | 50 | 29 | 29-Oct-2018 | 5.15 | 27565 |
| Bc5 | pole | 46 | 29 | 29-Oct-2018 | 19.36 | 25836 |
| B43 | pole | 48 | 26 | 31-Oct-2018 | 10.13 | 41416 |
| B44 | pole | 45 | 27 | 31-Oct-2018 | 22.41 | 21524 |
| BP5 | pole | 46 | 29 | 29-Oct-2018 | 96.23 | 29618 |
| B04 | pole | 46 | 29 | 29-Oct-2018 | 1.84 | 20708 |
| Pi2 | floor | 43 | 29 | 31-Oct-2018 | 16.88 | 74729 |
| Pi3 | floor | 38 | 28 | 31-Oct-2018 | 26.01 | 196087 |
| Pi4 | floor | 41 | 28 | 31-Oct-2018 | 23.92 | 75956 |
| Pi5 | floor | 41 | 27 | 31-Oct-2018 | 26.58 | 91192 |
| Pi1 | floor | 45 | 28 | 31-Oct-2018 | 127.80 | 63889 |
| AP1 | seat | 56 | 20 | 29-Oct-2018 | 22.19 | 46070 |
| AP2 | seat | 46 | 28 | 29-Oct-2018 | 2.65 | 64815 |
| AP3 | seat | 50 | 29 | 29-Oct-2018 | 2.77 | 94022 |
| AP4 | seat | 46 | 29 | 29-Oct-2018 | 1.45 | 53354 |
| AP5 | seat | 46 | 29 | 29-Oct-2018 | 17.05 | 26898 |
| AF2 | seat | 50 | 26 | 31-Oct-2018 | 2.63 | 26320 |
| AF3 | seat | 50 | 26 | 31-Oct-2018 | 21.47 | 30490 |
| AF1 | seat | 46 | 26 | 31-Oct-2018 | 18.55 | 68947 |
| AF4 | seat | 52 | 26 | 31-Oct-2018 | 1.62 | 39919 |
| AF5 | seat | 53 | 26 | 31-Oct-2018 | 2.46 | 75340 |

**Table S9.** Potentially pathogenic taxa detected.

| Taxa | Hands |  |  |  | Surfaces |  |  |  |  |  | sum |
| --- | --- | --- | --- | --- | --- | --- | --- | --- | --- | --- | --- |
|  | Before travel (not handwashing) (N=8) | After travel (not handwashing) (N=8) | Before travel (handwashing) (N=4) | After travel (handwashing) (N=8) | Poles (N=10) | Escalator handrails (N=5) | Platform floor (N=5) | Seats (N=10) | Stair handrails (N=5) | Turnstiles (N=5) |  |
| g__Streptococcus s__ | 8 | 8 | 4 | 8 | 10 | 5 | 5 | 10 | 5 | 5 | 68 |
| g__Staphylococcus s__epidermidis | 8 | 8 | 4 | 8 | 10 | 5 | 5 | 10 | 5 | 5 | 68 |
| g__Staphylococcus s__ | 8 | 8 | 4 | 8 | 10 | 5 | 5 | 10 | 5 | 5 | 68 |
| g__Propionibacterium s__acnes | 8 | 8 | 4 | 8 | 10 | 5 | 4 | 10 | 5 | 5 | 67 |
| g__Acinetobacter s__johnsonii | 7 | 8 | 3 | 8 | 9 | 5 | 5 | 10 | 4 | 5 | 64 |
| g__Pseudomonas s__stutzeri | 7 | 8 | 2 | 8 | 8 | 5 | 5 | 10 | 4 | 5 | 62 |
| g__Haemophilus s__parainfluenzae | 8 | 8 | 3 | 8 | 9 | 5 | 5 | 7 | 4 | 5 | 62 |
| g__Enterobacter s__hormaechei | 7 | 8 | 3 | 8 | 6 | 5 | 5 | 9 | 4 | 5 | 60 |
| g__Veillonella s__dispar | 7 | 8 | 2 | 8 | 7 | 5 | 5 | 6 | 4 | 5 | 57 |
| g__Rothia s__dentocariosa | 5 | 6 | 2 | 6 | 6 | 5 | 5 | 5 | 3 | 5 | 48 |
| g__Prevotella s__melaninogenica | 5 | 8 | 1 | 8 | 6 | 5 | 5 | 2 | 3 | 5 | 48 |
| g__Neisseria s__subflava | 6 | 7 | 1 | 6 | 7 | 5 | 4 | 4 | 3 | 4 | 47 |
| g__Propionibacterium s__granulosum | 5 | 8 | 3 | 7 | 8 | 5 | 0 | 2 | 3 | 4 | 45 |
| g__Rhodococcus s__fascians | 1 | 7 | 1 | 5 | 6 | 3 | 3 | 4 | 1 | 2 | 33 |
| g__Staphylococcus s__aureus | 2 | 5 | 0 | 4 | 5 | 5 | 4 | 4 | 1 | 3 | 33 |
| g__Veillonella s__parvula | 4 | 5 | 0 | 4 | 5 | 5 | 3 | 0 | 3 | 4 | 33 |
| g__Acinetobacter s__lwoffii | 3 | 5 | 0 | 2 | 3 | 3 | 4 | 6 | 3 | 1 | 30 |
| g__Staphylococcus s__saprophyticus | 2 | 5 | 1 | 3 | 3 | 4 | 1 | 5 | 0 | 4 | 28 |
| g__Streptococcus s__anginosus | 3 | 3 | 0 | 2 | 4 | 3 | 4 | 1 | 1 | 3 | 24 |
| g__Prevotella s__nigrescens | 1 | 5 | 0 | 3 | 2 | 5 | 3 | 0 | 1 | 2 | 22 |
| g__Porphyromonas s__endodontalis | 2 | 3 | 0 | 1 | 3 | 4 | 3 | 2 | 2 | 1 | 21 |
| g__Aeromonas s__hydrophila | 1 | 2 | 0 | 3 | 0 | 4 | 4 | 2 | 0 | 1 | 17 |
| g__Streptococcus s__sobrinus | 0 | 2 | 0 | 2 | 1 | 4 | 2 | 1 | 0 | 3 | 15 |
| g__Kingella s__denitrificans | 2 | 2 | 1 | 1 | 2 | 1 | 1 | 2 | 2 | 0 | 14 |
| g__Enterobacter s__gergoviae | 1 | 1 | 1 | 1 | 2 | 3 | 0 | 2 | 1 | 1 | 13 |
| g__Capnocytophaga s__ochracea | 1 | 3 | 0 | 2 | 0 | 2 | 1 | 1 | 1 | 2 | 13 |
| g__Neisseria s__cinerea | 1 | 2 | 0 | 3 | 0 | 3 | 1 | 2 | 1 | 0 | 13 |
| g__Aeromonas s__caviae | 0 | 3 | 1 | 2 | 0 | 2 | 1 | 2 | 0 | 1 | 12 |
| g__Prevotella s__intermedia | 0 | 4 | 0 | 2 | 1 | 2 | 1 | 0 | 2 | 0 | 12 |
| g__Pseudomonas s__pseudoalcaligenes | 0 | 1 | 0 | 3 | 0 | 2 | 2 | 1 | 1 | 0 | 10 |
| g__Streptococcus s__agalactiae | 1 | 0 | 0 | 1 | 1 | 2 | 1 | 0 | 1 | 1 | 8 |
| g__Selenomonas s__noxia | 1 | 1 | 0 | 0 | 1 | 2 | 0 | 0 | 1 | 2 | 8 |
| g__Serratia s__marcescens | 1 | 2 | 0 | 1 | 0 | 2 | 2 | 0 | 0 | 0 | 8 |
| g__Staphylococcus s__haemolyticus | 1 | 0 | 0 | 3 | 0 | 2 | 0 | 0 | 0 | 1 | 7 |
| g__Bacillus s__cereus | 1 | 0 | 0 | 0 | 0 | 3 | 0 | 1 | 0 | 2 | 7 |
| g__Pantoea s__agglomerans | 0 | 1 | 0 | 2 | 0 | 1 | 0 | 1 | 0 | 1 | 6 |
| g__Bacteroides s__uniformis | 0 | 1 | 0 | 2 | 0 | 1 | 0 | 0 | 1 | 1 | 6 |
| g__Pseudomonas s__alcaligenes | 0 | 1 | 0 | 1 | 0 | 0 | 3 | 1 | 0 | 0 | 6 |
| g__Haemophilus s__influenzae | 0 | 0 | 0 | 2 | 1 | 1 | 1 | 0 | 1 | 0 | 6 |
| g__Salmonella s__ | 1 | 1 | 0 | 1 | 0 | 1 | 0 | 0 | 0 | 1 | 5 |
| g__Bacteroides s__fragilis | 0 | 1 | 0 | 0 | 1 | 1 | 0 | 0 | 1 | 1 | 5 |
| g__Vibrio s__mimicus | 0 | 0 | 0 | 1 | 0 | 1 | 1 | 0 | 0 | 2 | 5 |
| g__Collinsella s__aerofaciens | 0 | 0 | 0 | 0 | 1 | 3 | 1 | 0 | 0 | 0 | 5 |
| g__Providencia s__stuartii | 0 | 1 | 0 | 0 | 0 | 1 | 1 | 0 | 0 | 1 | 4 |
| g__Ewingella s__americana | 0 | 1 | 0 | 1 | 0 | 0 | 1 | 1 | 0 | 0 | 4 |
| g__Streptobacillus s__moniliformis | 2 | 0 | 0 | 1 | 0 | 0 | 0 | 0 | 0 | 0 | 3 |
| g__Plesiomonas s__shigelloides | 0 | 0 | 0 | 0 | 0 | 1 | 1 | 0 | 1 | 0 | 3 |
| g__Brevundimonas s__diminuta | 0 | 1 | 0 | 0 | 0 | 1 | 0 | 0 | 1 | 0 | 3 |
| g__Mycobacterium s__celatum | 0 | 0 | 0 | 0 | 0 | 0 | 0 | 1 | 0 | 1 | 2 |
| g__Listeria s__seeligeri | 0 | 0 | 0 | 1 | 0 | 0 | 1 | 0 | 0 | 0 | 2 |
| g__Enterococcus s__casseliflavus | 1 | 0 | 0 | 1 | 0 | 0 | 0 | 0 | 0 | 0 | 2 |
| g__Clostridium s__perfringens | 0 | 1 | 0 | 1 | 0 | 0 | 0 | 0 | 0 | 0 | 2 |
| g__Bacillus s__coagulans | 0 | 2 | 0 | 0 | 0 | 0 | 0 | 0 | 0 | 0 | 2 |
| g__Rhodococcus s__equi | 0 | 0 | 0 | 0 | 1 | 0 | 0 | 0 | 0 | 0 | 1 |
| g__Megamonas s__hypermegale | 0 | 1 | 0 | 0 | 0 | 0 | 0 | 0 | 0 | 0 | 1 |
| g__Legionella s__pneumophila | 0 | 0 | 0 | 0 | 0 | 1 | 0 | 0 | 0 | 0 | 1 |
| g__Capnocytophaga s__canimorsus | 1 | 0 | 0 | 0 | 0 | 0 | 0 | 0 | 0 | 0 | 1 |
| g__Bacteroides s__caccae | 0 | 1 | 0 | 0 | 0 | 0 | 0 | 0 | 0 | 0 | 1 |
| g__Moraxella s__lincolnii | 0 | 0 | 0 | 0 | 0 | 0 | 0 | 0 | 0 | 1 | 1 |

